# Foraging for locally and spatially varying resources: Where exploitation competition, local adaptation and kin selection meet

**DOI:** 10.1101/2022.10.03.510600

**Authors:** Max Schmid, Claus Rueffler, Laurent Lehmann, Charles Mullon

## Abstract

In patch- or habitat-structured populations different processes can lead to diversity at different scales. While spatial heterogeneity generates spatially disruptive selection favoring variation between patches, local competition can lead to locally disruptive selection promoting variation within patches. So far, almost all theory has studied these two processes in isolation. Here, we use mathematical modelling to investigate how resource variation within and between habitats influences the evolution of variation in a consumer population where individuals compete in finite patches connected by dispersal. We find that locally and spatially disruptive selection typically act in concert, favoring polymorphism under a significantly wider range of conditions than when in isolation. But when patches are small and dispersal between them is low, kin competition inhibits the emergence of polymorphism, especially when driven by local competition. We further use our model to clarify what comparisons between trait and neutral genetic differentiation (*Q*_st_/*F*_st_ comparisons) can tell about the nature of selection. Overall, our results help understand the interaction between two major drivers of diversity: locally and spatially disruptive selection; and how this interaction is modulated by the unavoidable effects of kin selection under limited dispersal.

## 1 Introduction

Adaptation to exploit different resources has long been recognized as one of the major drivers of phenotypic diversity (Skúlason and Smith, 1995; Smith and Skúlason, 1996). One example of such resource-driven diversity can be found in the three-spined stickleback, *Gasterosteus aculeatus*, which has diverged between lake and stream habitats in multiple locations. Across locations, lake sticklebacks have additional and longer gill rakers allowing them to capture zooplankton, whereas stream sticklebacks have fewer and shorter rakers to feed on benthic macroinvertebrats (Hendry et al., 2002; Berner et al., 2010; Ravinet et al., 2013). Another emblematic example comes from Darwin’s ground finches, *Geospiza*. Within the same island, different species of this genus show highly-diverged beak morphologies, with each form fitting to a specific resource: large-beaked finches are specialized to large and hard seeds while small pointed-beaked finches are specialized to smaller and softer food sources (Grant and Grant, 2002, 2008).

The examples of three-spined stickleback and Darwin’s finches in fact each illustrate one main pathway that has been proposed to lead to resource polymorphism. Where resources vary between habitats, for instance when lakes and streams offer different diets, diversity is driven by *spatially disruptive selection* as different traits are favored in different locations. In this case, polymorphism leads to local adaptation where each morph shows a better fit to the habitat it lives in (Haldane, 1948; Kawecki and Ebert, 2004; Leimu and Fischer, 2008; Hereford, 2009). In contrast, when resources vary within habitats, for instance when each island offers a wide range of seeds, diversification is driven by local competition. This leads to character displacement owing to negative frequency-dependent selection as individuals feeding on food sources alternative to others enjoy less intense competition (Maynard Smith, 1962; Rosenzweig, 1978; Dieckmann and Doebeli, 1999; Rueffler et al., 2006*a*). To contrast with spatially disruptive selection, we will refer to such selection favoring polymorphism due to local competition as *locally disruptive selection*.

Mathematical models have been useful to understand how spatially (e.g. Levene, 1953; Felsenstein, 1976; Brown and Pavlovic, 1992; Meszéna et al., 1997; Geritz and Kisdi, 2000; Svardal et al., 2015) and locally (e.g. MacArthur and Levins, 1967; Roughgarden, 1976; Christiansen and Loeschcke, 1980; Slatkin, 1980; Abrams, 1986; Meszéna et al., 1997; Geritz et al., 1998; Day, 2000, 2001; Ajar, 2003; Rueffler et al., 2006*b*; Abrams et al., 2008) disruptive selection can lead to trait diversity within and between species. One salient point from these models is the antagonistic effects of dispersal and gene flow between habitats on polymorphism. On the one hand, limited dispersal favors the emergence of polymorphism under spatially disruptive selection as it allows different morphs to become associated to different habitats (Levene, 1953; Felsenstein, 1976; Lenormand, 2002). On the other hand, limited dispersal inhibits polymorphism when driven by locally disruptive selection because it leads to interactions among kin who are not sufficiently diverged to escape local competition – i.e., kin selection opposes polymorphism (Day, 2001; Ajar, 2003).

Current models of resource polymorphism have focused exclusively on either spatially or locally disruptive selection (citations in preceding paragraph), in effect assuming that resources vary only between or only within patches (though see Day, 2000 for an analysis ignoring kin selection). More realistically, however, variation may occur both between and within patches leading to both selection forces acting simultaneously. Given the antagonistic effects of dispersal on spatially and locally disruptive selection, the outcome of evolutionary dynamics in this case is unclear. To investigate this, we model the evolution of a consumer trait when resources vary within and between patches of finite size that are connected by limited dispersal. Our results suggest that in fact spatially and locally disruptive selection may very often work together to promote the emergence and maintenance of resource polymorphism.

## 2 The model

### 2.1 Population and life cycle events

We consider an asexual population that is divided among a large (ideally infinite) number of heterogeneous patches and that is censused at discrete demographic time points (henceforth referred to as “years” but the model applies to any time period that the species under consideration needs to complete its life cycle). At the beginning of each year, all patches carry the same number *n* of adult individuals but differ according to the resources they hold (we detail this in the next section 2.2). The following events then unfold within a year, determining the life cycle (Fig. 1a): (1) resource consumption and reproduction: within each patch, adults first consume local resources and then reproduce clonally, making a large number of offspring (how consumption is modeled is specified in the next section 2.2); (2) dispersal: each offspring either remains in its natal patch (with probability 1−*m*) or disperses to another randomly sampled patch (with probability *m*); (3) survival: each adult survives or dies (with probabilities *γ* and 1−*γ*, respectively), in the latter case freeing up a breeding spot or territory within its patch; and finally (4) population regulation: philopatric and immigrant offspring compete locally for open spots to become adults, so that by the end of the year, each patch again carries *n* adult individuals.

**Figure 1:**
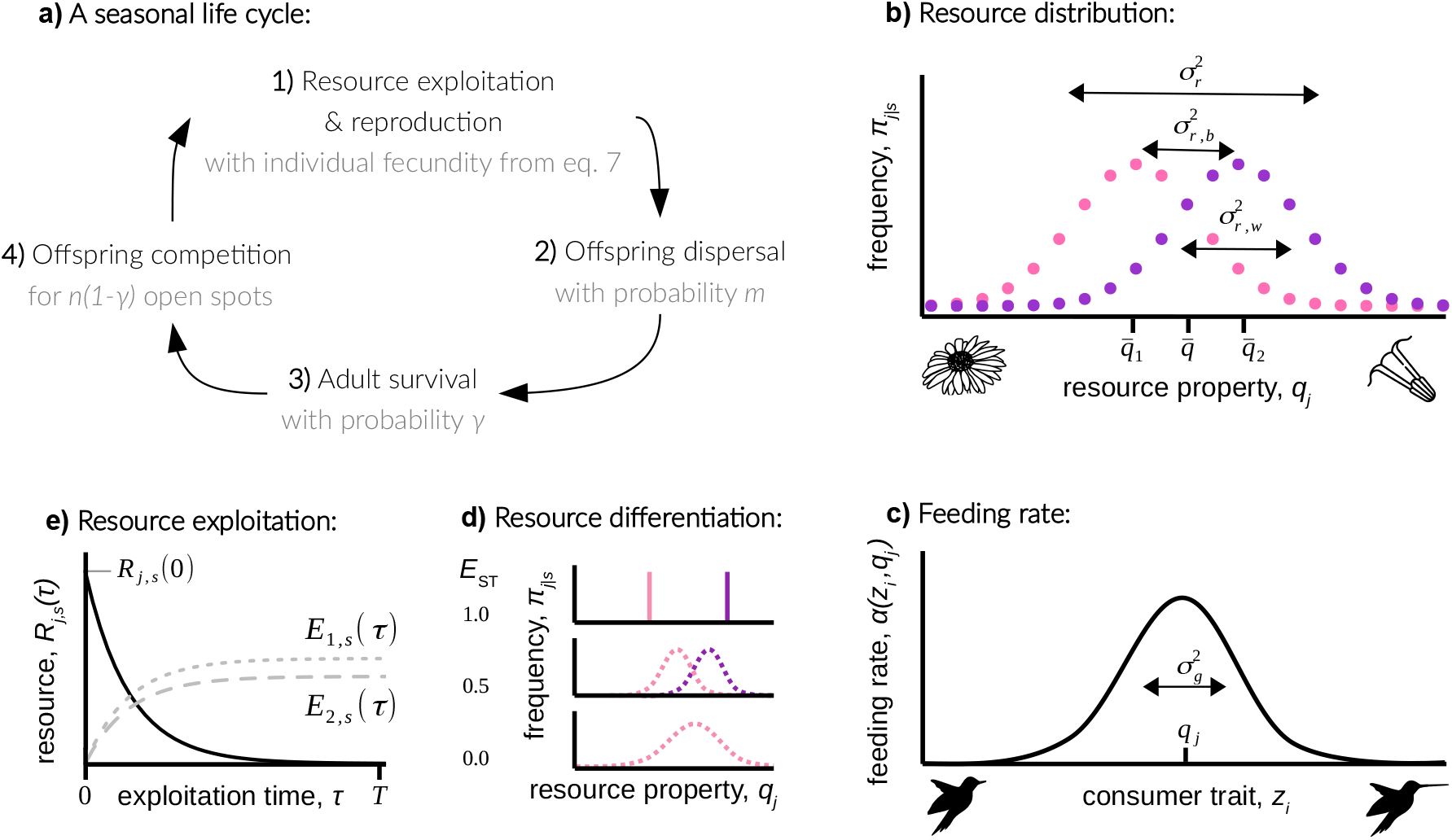
A model of resource exploitation for a spatially-structured population. **a)** Sequence of life-cycle events within each year (section 2.1 for details). **b)** Resource distribution *π*_*j*|*s*_ within patches of two types *s =* 1 (pink) and *s =* 2 (purple). Resources vary both within 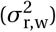 and between 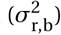 patches (section 2.2.1, eq. 1). **c)** Feeding rate *α*(*z*_*i*_, *q*_*j*_) of individual *i* on resources of type *j* with property *q*_*j*_ as a function of consumer trait *z*_*i*_ (eq. 3), which is maximal when the trait matches the resource property, i.e. *z*_*i*_ *= q*_*j*_. **d)** Resource differentiation among patches *E*_ST_ (eq. 2) for different resource distributions *π*_*j*|*s*_ within patches of two types *s =* 1 (pink) and *s =* 2 (purple). When *E*_ST_ *=* 1, each habitat contains a single different resource; when *E*_ST_ *=* 0, there is a single habitat offering a range of resources. **e)** Within-year resource dynamics according to eq. (4) with the amount *R*_*j, s*_ (*τ*) of resource *j* in a patch of type *s* in black and the energy *E*_1,*s*_ (*τ*) (in gray dotted) and *E*_2,*s*_ (*τ*) (in gray dashed) accumulated by two individuals where individual 1 expresses a trait that allows to extract more resources at the expense of individual 2.

### 2.2 Resource distribution and consumption

#### 2.2.1 Ecological variation within and between patches

We assume that individuals consume a resource that varies in some relevant quantitative property within and between patches (e.g., corolla length, prey speed). To describe this variation, we let 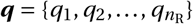 denote the set of possible values the resource can take where *q*_*j*_ ∈ ℝ is the value of the *j* -th resource type (for instance, if ***q*** *=* {2, 4}, then the resource can take the value *q*_1_ *=* 2 and *q*_2_ *=* 4). Before consumption, a patch is then characterized by a frequency distribution over ***q*** (for instance, the frequency of resource of type *q*_1_ *=* 2 is 0.2 and that of type *q*_2_ *=* 4 is 0.8) and we assume that there is a finite number of such possible frequency distributions. Specifically, let Π_*s*_ *=* {*π*_1|*s*_, *π*_2|*s*_, …} stand for the *s*-th such frequency distribution over the resource values where *π*_*j* |*s*_ is the frequency of a resource of type *j* in a patch characterized by this *s*-th frequency distribution. We say that a patch is in state *s* ∈ Ω if its resources distribution is Π_*s*_ and denote by *π*_*s*_ the frequency of patches in state *s* in the population (i.e., ∑_*s*∈Ω_ *π*_*s*_ *=* 1). The set of patches that are in the same state *s* belong to the same habitat.

A patch in state *s* thus has a resource with average property 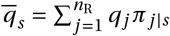, and variance 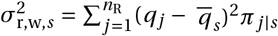. At the global level, the resource has an average property 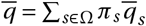 and variance

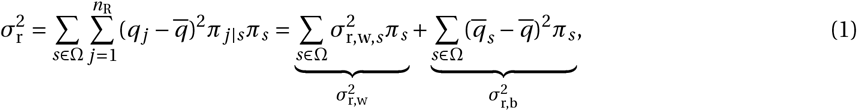

which we decomposed into the average within-patch resource variance 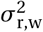 and the variance across patches 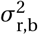 (Fig. 1b). This decomposition allows us to quantify the level of resource differentiation across patches with

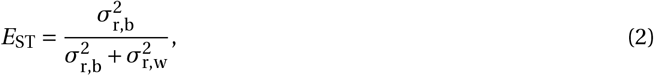

which denotes the proportion of variance that is due to variance between patches (Fig. 1d). When *E*_ST_ *=* 1, all resource variation is between patches (Fig. 1d, top), and only within patches when *E*_ST_ *=* 0 (Fig. 1d, bottom).

#### 2.2.2 Trait-based competition for resources

Each year, individuals consume local resources within their patch (step (1) in section 2.1). We assume that their ability to consume these resources depends on a quantitative trait they express. For instance, the ability of a hummingbird to extract nectar depends on how well the length of its bill matches the length of the corolla tube of the flowers it visits. To capture such a situation or, more generally, a scenario of trait-based resource consumption, we let the rate at which an individual indexed as *i* ∈ {1, 2,…, *n*} in a focal patch (recall that *n* is the number of adults in a patch) feeds on a resource of type *j* be

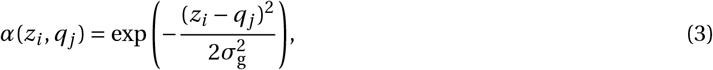

where *z*_*i*_ ∈ ℝ is the value of a relevant trait expressed by this individual. According to eq. (3), the feeding rate of an individual on resource *j* is maximal when its trait matches the resource property, *z*_*i*_ *= q*_*j*_, and declines with increasing distance between *z*_*i*_ and *q*_*j*_, with the rate of decline inversely related to 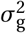 (Fig. 1c). The parameter 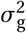 thus describes to what extent individuals can be generalists: an individual with trait *z*_*i*_ can feed at a high rate on wider range of resource types when 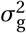 is large compared to when 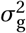 is small.

We assume that adults feed over a time interval of length *T* (taking place within step (1) of the year, see section 2.1). The variable *T* thus controls how much time individuals have to consume resources within a year.

If we denote by *R*_*j, s*_ (*τ*) the density of the resource *j* in a patch of type *s* at time *τ* (0 ≤ *τ* ≤ *T*) and by *E*_*i, s*_ (*τ*) the energy accumulated by individual *i* via consumption in that patch at time *τ*, then these variables change according to

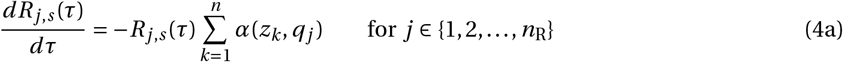

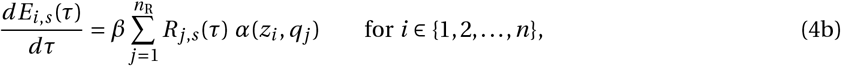

where *β* is the energy content per resource unit. Individuals with a higher feeding rate *α*(*z*_*i*_, *q*_*j*_) thus accumulate more resources of type *j* at the expense of other individuals in the patch (Fig. 1e). The quantity of a resource *j* at the beginning of each year is determined by the state *s* of the patch, i.e., by

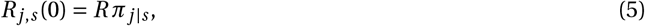

where *R* is the total density of resources, which is assumed to be the same in all patches. Resources are thus self-renewing and brought back to their common carrying capacity *R* each year. The energy budget of each individual is initially zero, i.e. *E*_*i, s*_ (0) *=* 0. Eq. (4) can then be solved to obtain the net energy uptake *E*_*i, s*_ (*T*) that an individual *i* has accumulated by time *T*. In fact, the net energy uptake of individual *i* in a focal patch of type *s* obtained from resource *j* can be written as

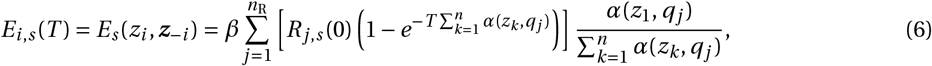

which depends on the trait *z*_*i*_ of the individual and the collection ***z***_−*i*_ *=* (*z*_1_, *z*_2_,…, *z*_*i* −1_, *z*_*i +*1_,…, *z*_*n*_) of traits expressed by its *n* − 1 patch neighbors (Appendix A.1, eq. A1). The term within square brackets in eq. (6) corresponds to the total amount of resource *j* that is consumed within a time period in a focal patch that is in state *s*. As the exploitation time increases (i.e., as *T* → ∞), this converges to the amount of resources available, *R*_*j, s*_ (0), as individuals have sufficient time to consume all resources in a patch. The ratio in eq. (6), meanwhile, gives the share of these resources that are collected by the focal individual (which simplifies to (1/*n*) in a monomorphic population, i.e., when *z*_*k*_ *= z* for all *k*). As such, eq. (6) in the limit *T* → ∞ corresponds to a contest success function of the ratio type, which is commonly used to model conflict over resources (Hirshleifer, 1989). Our dynamical model eq. (4) thus provides a mechanistic derivation for such a contest success function.

### 2.3 Fecundity, fitness, and evolutionary dynamics

We assume that the fecundity of an individual (the total number of offspring it produces during step (2) of the life-cycle) is proportional to the total energy it has accumulated, i.e., the fecundity of individual *i* with trait *z*_*i*_ in a patch in state *s* and where its neighbors have traits ***z***_−*i*_ is given by

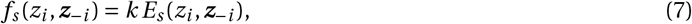

where *k* is the conversion factor from energy to offspring. From this fecundity, we can characterize an individual’s fitness, which here is defined as the expected number of successful offspring produced by this individual over one full year (those that establish as adults, including itself if it survives). This individual fitness measure, which we detail in Appendix A.2, lays the basis of our evolutionary analysis.

We are interested in the genetic evolution of the consumer trait *z*, in particular, whether gradual evolution can result in polymorphism. Our approach is based on Ohtsuki et al. (2020) (see our appendix B for details). We assume that mutations are rare with small phenotypic effect such that evolutionary dynamics occur in two steps. First, the population evolves gradually under directional selection via the constant input of mutations. The population eventually converges to a singular phenotype, which we denote as *z*^*^, where directional selection ceases to act. Once the population expresses *z*^*^, the population either experiences stabilising selection and remains monomorphic, or experiences disruptive selection and becomes polymorphic. The process of transitioning from a monomorphic to a dimorphic phenotype distribution is referred to as “evolutionary branching” (Geritz et al., 1998). We determine the conditions under which polymorphism emerges in our model in Appendix C and summarise our main findings below. Beyond the specific model of resource competition investigated here, our analysis of selection as described in appendix B can in fact accommodate for any trait that influences fecundity in group-structured populations (under our life-cycle assumptions given in section 2.1). This analysis therefore allows to understand the emergence of polymorphism in a wide range of social traits in terms of their fecundity effects in populations where limited dispersal leads to interactions among relatives (see in particular eqs. B10 and eqs. B24-B25 in appendix B).

## 3 Results

### 3.1 How resource variation between and within patches favors polymorphism

We first assume that exploitation time is long (i.e., *T* → ∞) as this allows for an in-depth mathematical analysis. In this scenario, all available resources are consumed by the end of the exploitation period in each patch, and, as a result, the same total number of offspring is produced in each patch (so that selection is “soft”, Wallace, 1975; Christiansen, 1975; Débarre and Gandon, 2011). For this case, we find that the population first evolves toward the singular trait value

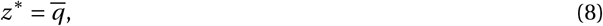

where the consumer trait matches the average resource property (appendix C.1.2 for derivation; in line with models based on soft selection, e.g., Geritz et al., 1998; Day, 2001; Ajar, 2003; Svardal et al., 2015; Ohtsuki et al., 2020). Once the population has converged to express 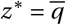, our analysis (appendix C.1.3) shows that selection is disruptive and therefore favors the emergence of polymorphism when

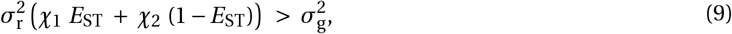

where

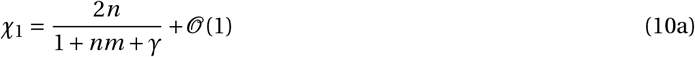

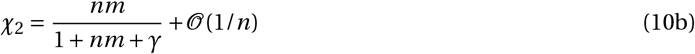

are complicated non-negative quantities that depend on demographic parameters but whose remainders vanish in the limit of large patch size and low dispersal (i.e., when *n* → ∞ and *m* → 0 such that *nm* remains constant; see eqs. C28 & C30 in the Appendix for the full expressions).

Condition (9) reveals that the evolution of polymorphism depends on several factors. First, polymorphism is favored when the total resource variance is large compared to the degree of consumer generalism (when 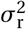 is large and 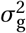 is small). This is easy to intuit and reflects that greater resource diversity 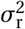 provides more ecological opportunities and facilitates the coexistence of specialized consumers. In contrast, a generalist consumer with large 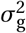 can successfully consume a wide range of resources and thereby prevent the emergence of specialized morphs.

The total variation in resources 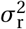 can be partitioned into variation between and within patches (according to *E*_ST_ and 1−*E*_ST_ respectively, see eqs. 1-2). Eq. (9) reveals that both types of variation favor evolutionary branching. Specifically, these two sources of variation add up to promote polymorphism, with variation between patches weighted by *χ*_1_ and within patches by *χ*_2_. Between-patch variation thus contributes most to disruptive selection when *χ*_1_*E*_ST_ is large compared to *χ*_2_(1−*E*_ST_). If this is so and condition (9) holds, polymorphism is primarily driven by spatially disruptive selection favoring local adaptation, i.e., favoring individuals whose traits match their local resource average. Inspection of *χ*_1_ shows that it is most sensitive to dispersal *m*, rapidly decreasing as dispersal increases (eq. 10a; Fig. 3a). This reflects the well-known notion that gene flow inhibits local adaptation as it homogenizes genetic variation between patches (e.g. Day, 2000; Lenormand, 2002). Furthermore, *χ*_1_ decreases as adult survival *γ* increases and patch size *n* decreases (eq. 10a; Fig. 3a; Fig. S1), indicating that spatially disruptive selection is weaker when generation overlap and patches are small. This is because these conditions lead to stronger kin competition, which reduces the strength of selection on competitive traits as an individual’s reproductive success comes at the expense of relatives.

**Figure 2:**
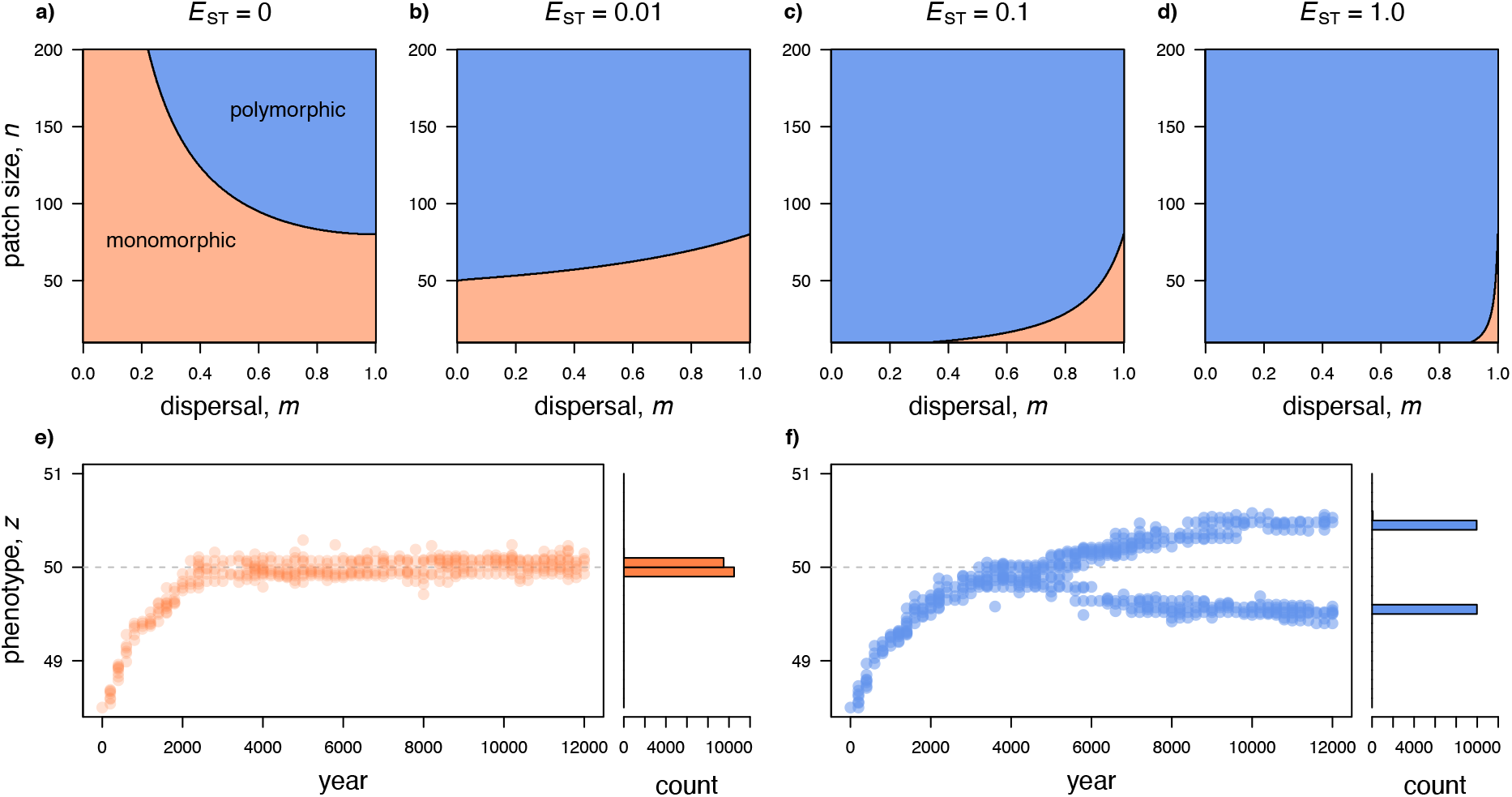
The evolution of polymorphism with locally and spatially varying resources. **a)-d)** Parameter space in which selection is stabilising (pink) or disruptive (blue) from eq. (9) for four levels of resource differentiation: a) *E*_ST_ *=* 0; b) *E*_ST_ *=* 0.01; c) *E*_ST_ *=* 0.1; d) *E*_ST_ *=* 1 (other parameters 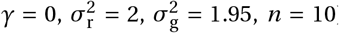). **e)-f)** Simulated evolution of the consumer trait *z* when selection is e) stabilising (*E*_ST_ *=* 0 and 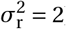) and f) disruptive (*E*_ST_ *=* 1 and 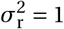). See appendix D for details on the simulation procedure. Other parameters: *n =* 10, *γ =* 0, *m =* 0.1, and 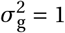. Each segregating phenotype every 50 years is represented by a filled circle. Gray dashed line indicates the singular trait value: *z*^*^ *=* 50. As predicted by our mathematical analysis, the population first converges to *z*^*^, and then remains monomorphic when selection is stabilising (e) or becomes polymorphic when selection is disruptive (f).

**Figure 3:**
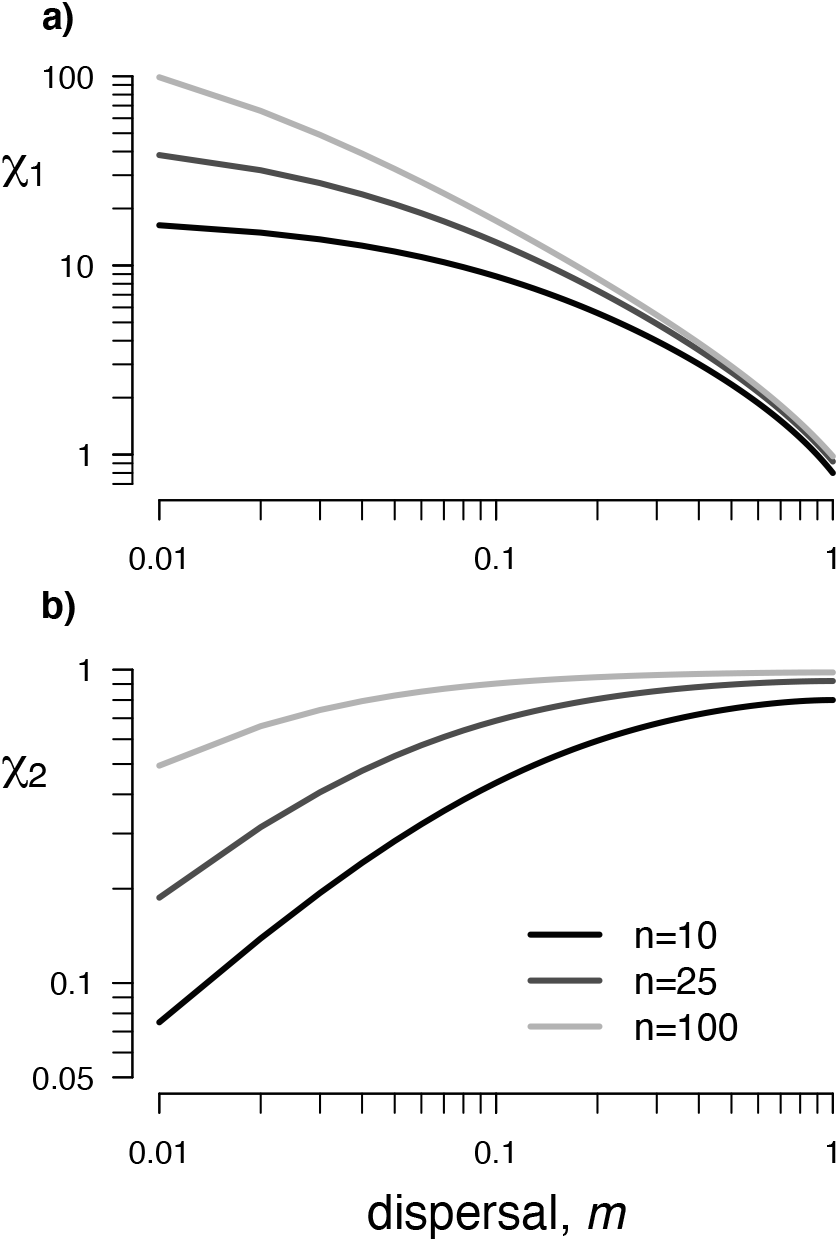
Disruptive selection and the weights for spatial and local resource variation in eq. 9. **a)** *χ*_1_ and **b)** *χ*_2_ with *n =* 10 (black), 25 (dark gray), and 100 (light gray; from eq. C28-C30 in Appendix; other parameters: *γ =* 0). In a panmictic population (i.e. when *m =* 1), both weights are equal, *χ*_1_ *= χ*_2_ *=* 1−2/*n*. Otherwise, *χ*_1_ *> χ*_2_ indicating that spatial resource variation typically plays a bigger role than local variation. While *χ*_1_ decreases, *χ*_2_ increases with dispersal, showing that spatially and locally disruptive selection are favored and disfavored, respectively, by limited dispersal.

Conversely, within-patch resource variation contributes most to disruptive selection when *χ*_2_(1−*E*_ST_) is large compared to *χ*_1_*E*_ST_. If so and condition (9) holds, polymorphism is primarily driven by locally disruptive selection, or, more specifically, negative frequency-dependent disruptive selection at the local scale. This selection favors rare morphs in each patch because these morphs are able to exploit resources that are under less intense competition. In contrast to *χ*_1_, the weighting factor *χ*_2_ increases with dispersal *m* (eq. 10b; Fig. 3b; Fig. S1 for effect of adult survival *γ* and patch size *n*). This is because when dispersal is weak, individuals in the same patch tend to express similar traits as they are more genetically related and as a result cannot escape local competition (Day, 2001; Ajar, 2003).

Our condition (9) is in perfect agreement with previous expressions derived for less general ecological models. In particular, when resources vary only within patches (*E*_ST_ *=* 0) and generations do not overlap (*γ =* 0), we recover the polymorphism condition of Ajar (2003) (his eq. 25). When resources vary only between patches (*E*_ST_ *=* 1), our inequality (9) reduces to eq. (66) of Ohtsuki et al. (2020) under high adult survival (i.e., *γ* ∼ 1, akin to a Moran model), to eq. (C.15) of Svardal et al. (2015) under no adult survival and infinite patch size (*γ =* 0 and *n* → ∞), and to eq. (B.8) of Geritz et al. (1998) when additionally assuming panmixia (*m =* 1). In contrast to these previous studies, our result holds for arbitrary resource distributions both within and between habitats (in addition to allowing for limited dispersal, finite patch size and intermediate adult survival). This reveals that dispersal *m* can either hinder or favor the emergence of polymorphism, depending on whether resource variation is primarily distributed between or within patches. However, even when between-patch differentiation *E*_ST_ *>* 0 is relatively low, dispersal limitation favors rather than hinders polymorphism (Fig. 2b-c). This boils down to the fact that *χ*_1_ ≥ *χ*_2_ in eq. (9) (with equality only when *m =* 1, see eq. C31 and fig. S1a). Thus, under limited dispersal spatially disruptive selection due to between-patch variation has a greater weight than locally disruptive selection due to within-patch variation in determining whether polymorphism emerges. In other words, polymorphism is more likely to be due to local adaptation than local competition when gene flow is limited (Fig. 2a-d).

### 3.2 Gradual emergence of polymorphism, its distribution and genetic signatures

Condition (9) determines whether or not selection favors the emergence of polymorphism. But it does not inform us about the long term fate of the morphs, specifically, their final trait values and their distribution within and between patches. To investigate this, we ran stochastic individual-based simulations for a population occupying 2000 patches (using Nemo-Age, Cotto et al., 2020; see Appendix D for details). We explored various combinations of parameters, in particular allowing for various distributions of resources within and between patches (by varying *E*_ST_). As predicted by our analysis, the population first converges to match the average resource property 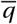 while remaining largely monomorphic (Fig. 2e,f). When eq. (9) is not satisfied, the population remains monomorphic with a unimodal phenotypic distribution (Fig. 4a,b in pink). But when eq. (9) is satisfied, the population becomes polymorphic with two highly differentiated morphs eventually coexisting in the population: one that specializes on “small” resources (small *q*-values) and the other on “large” resources (large *q*-values, Fig. 2e,f; Fig. 4a,b in blue).

**Figure 4:**
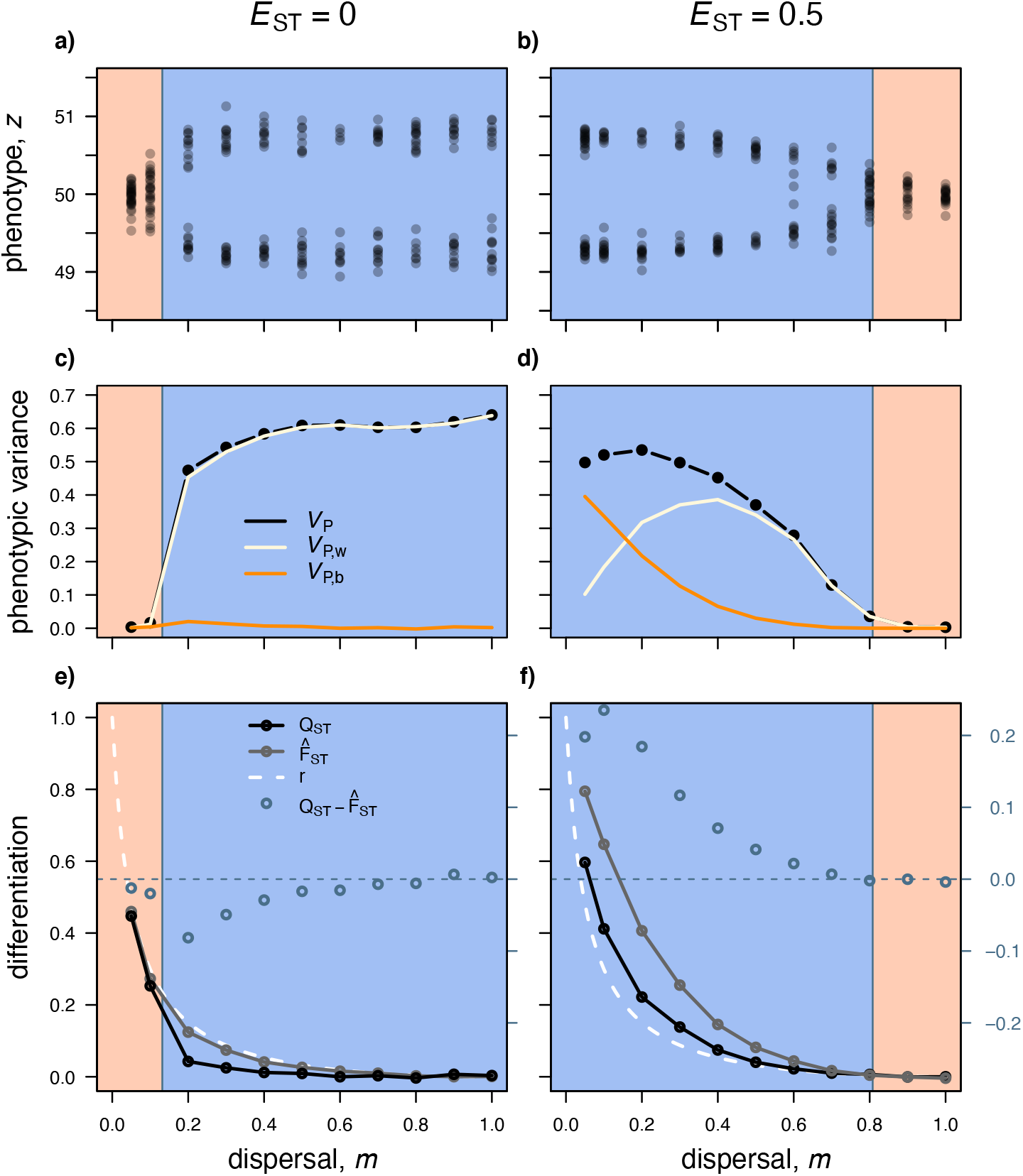
The distribution of polymorphism within and between patches. **a)-b)** Unique trait values in a simulated population after 2,000 years of evolution as a function of dispersal *m* with a) *E*_ST_ *=* 0 and b) *E*_ST_ *=* 0.5 (other parameters: 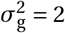 with *E*_ST_ *=* 0, 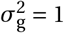 with *E*_ST_ *=* 0.5, *γ =* 0, and *n =* 10). Blue and pink areas indicate levels of dispersal that lead to disruptive and stabilising selection, respectively (from eq. 9). Simulations and mathematical analyses are thus consistent, with the population evolving to being dimorphic where selection is disruptive (blue) and monomorphic where selection is stabilising (pink). **c)-d)** Phenotypic variance in the same evolved populations as in a) and b) with total variance *V*_P_ in black, decomposed as variance within patches *V*_P,w_ (white) and between patches *V*_P,b_ (orange). Phenotypic variance is thus significantly greater when selection is disruptive (blue area). **e)-f)** Phenotypic differentiation *Q*_ST_ (in black) vs. neutral genetic differentiation quantified with 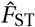 (in gray) from simulations (using the Weir-Cockerham approach with *hierfstat* package in R) and pairwise relatedness *r* (dashed white) from our mathematical model (using eq. B8). Dark blue open circles give 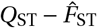 with scale given on the right hand side, blue dashed line indicates 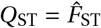. This shows that 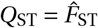 when selection is stabilising (pink), 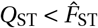 when selection is locally disruptive (i.e. when *E*_ST_ *=* 0, blue area in a), and 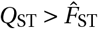 when selection is disruptive and *E*_ST_ *>* 0 (blue area in b).

The emergence of polymorphism is typically accompanied by a significant increase in population phenotypic variance (denoted as *V*_P_, Fig. 4c,d in black). The final level of phenotypic variance *V*_P_ maintained in a polymor-phic population depends on gene flow and how resources are distributed. When resource variation is mostly within patches (i.e., small *E*_ST_), *V*_P_ increases with dispersal (Fig. 4c). When resources vary primarily between patches (i.e., large *E*_ST_), variance *V*_P_ decreases with dispersal (Fig. 4d). This reflects how gene flow interacts differently with locally and spatially disruptive selection (as elaborated in section 3.1). In addition, phenotypic variance increases with *E*_ST_ (Fig. S3). In other words, two morphs maintained by spatially disruptive selection tend to be differentiated more strongly than two morphs maintained by purely locally disruptive selection. This further supports the notion that spatially disruptive selection is a stronger driver of polymorphism than locally disruptive selection.

To explore how the two morphs are distributed among patches, it is useful to decompose the phenotypic variance into within (*V*_P,w_) and between (*V*_P,b_) patch trait variance, *V*_P_ *= V*_P,w_ *+ V*_P,b_. As expected, within-patch phenotypic variance *V*_P,w_ is greatest when resource variation occurs within patches and dispersal is high (i.e., small *E*_ST_ and large *m*, Fig. 4c in white), while between-patch phenotypic variance *V*_P,b_ dominates when resource variation is concentrated among patches and dispersal is limited (i.e., large *E*_ST_ and small *m*, Fig. 4d in orange). This indicates that both morphs tend to co-occur in the same patch in the former case, while the different morphs tend to inhabit different patches in the latter case. Interestingly, as long as *E*_ST_ *>* 0, within-patch phenotypic variance responds non-monotonically to gene flow, with *V*_P,w_ initially increasing but ultimately decreasing with dispersal *m* (Fig. 4d in white). This pattern can be understood by considering that when resources are differentiated between patches (*E*_ST_ *>* 0) and patches are isolated (*m =* 0), patches tend to be fixed for different phenotypes so that *V*_P,w_ *=* 0. As dispersal increases, these different phenotypes start mixing within patches leading initially to an increase in within-patch variance *V*_P,w_. But past a threshold of dispersal, gene flow counteracts differentiation among patches, which eventually generates a decline in *V*_P,w_ (as reported in simulation studies with *E*_ST_ *=* 1, e.g. McDonald and Yeaman, 2018).

How gene flow and selection shape the distribution of morphs among patches can be further investigated through *Q*_ST_ *= V*_G,b_/(*V*_G,b_ *+ V*_G,w_), which measures among-patch differentiation in additive genetic variance (which equals phenotypic differentiation in the absence of environmental effects on the phenotype so that *V*_G,b_ *= V*_P,b_ and *V*_G,w_ *= V*_P,w_, like in our model). In particular, comparing *Q*_ST_ with genetic differentiation 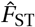 at neutral loci allows to focus on the effects of selection on phenotypic differentiation (e.g., Whitlock, 2008; Ovaskainen et al., 2011; Leinonen et al., 2013). We compute 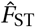 from our simulations, where in addition to a single locus coding for the consumer trait *z*, individuals carry a neutral locus (similar to a microsatellite marker) at which we compute genetic differentiation following the Weir-Cockerham approach (Weir and Cockerham, 1984; Appendix D for details). We also quantify genetic differentiation from our analytical model via pairwise relatedness *r*, i.e., the probability that two individuals randomly sampled without replacement from the same patch are identical-by-descent in a population monomorphic for the singular strategy, 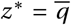 (see eq. B8 in appendix B.2.3).

As previously demonstrated, the two measures of genetic differentiation, *r* and 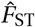, match perfectly in monomorphic populations (when eq. 9 does not hold, Rousset, 2002; Fig. 4e,f pink region) and in polymorphic populations where *E*_ST_ *=* 0 (Mullon and Lehmann, 2019; Fig. 4e blue region). However, in the presence of polymorphism (eq. 9 holds) and ecological differentiation among patches (*E*_ST_ *>* 0), 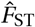 is greater than *r* (Fig. 3f blue region). The reason is that effective gene flow is diminished in a population consisting of two coexisting morphs that are adapted to local patch conditions. While 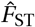 from our simulations detects this reduction in gene flow, the analytical neutral relatedness measure *r* cannot (as it is based on a monomorphic population).

Comparing phenotypic differentiation *Q*_ST_ with neutral genetic differentiation (quantified either with 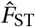 from a neutral marker in our simulations or relatedness *r* from the analytical model) reveals three patterns (Fig. 4e,f blue circles): (i) In the absence of polymorphism, *Q*_ST_ equals neutral genetic differentiation (Fig. 4e,f pink regions). (ii) In the presence of polymorphism and resource variation among patches (*E*_ST_ *>* 0), *Q*_ST_ exceeds neutral genetic differentiation (Fig. 4f blue region). (iii) In the presence of polymorphism and resource variation only within patches (*E*_ST_ *=* 0), *Q*_ST_ is lower than neutral genetic differentiation (Fig. 4e blue region). In other words, when the trait is under stabilizing selection for the same value in all patches so that the population is monomorphic, trait differentiation *Q*_ST_ is identical to neutral 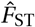. But when resource variation leads to polymorphism, *Q*_ST_ deviates from 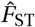. Spatially disruptive selection leads to an increase in between-patch phenotypic variance *V*_P,b_ and therefore to an increase in *Q*_ST_ relative to 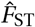. Conversely, locally disruptive selection boosts within-patch phenotypic variance *V*_P,w_, causing a drop in *Q*_ST_ compared to 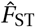.

Our observation that phenotypic differentiation exceeds neutral genetic differentiation 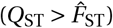 in the presence of polymorphism and resource differentiation (*E*_ST_ *>* 0) aligns with the common notion that 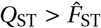 signals local adaptation (e.g. Whitlock, 2008; Leinonen et al., 2013). Another common idea is that 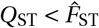 is an indicator of stabilising selection favoring the same trait value in all patches (i.e., under spatially uniform selection where *E*_ST_ *=* 0 and eq. 9 does not hold, e.g. Merilä and Crnokrak, 2001; McKay and Latta, 2002; Leinonen et al., 2008). By contrast, we find that phenotypic differentiation is actually similar to neutral genetic differentiation in this case, 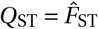 (this has also been demonstrated mathematically in Mullon and Lehmann, 2019, their eq. C26). Rather, we find 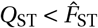 when selection is disruptive due to local competition only (eq. 9 holds and *E*_ST_ *=* 0) as this maintains greater phenotypic variation within patches than expected under neutrality (as suggested by Lamy et al., 2012).

### 3.3 Hard selection and alternative routes to polymorphism

Our results so far are based on the assumption that exploitation time *T* is long such that all resources are consumed within each year, resulting in the total number of offspring born in each patch to be the same (i.e. leading to soft selection). To relax this, we assume here that exploitation time *T* is short such that individuals do not interfere with one another through resource consumption (eq. C32 in Appendix C.2.1 for details; Figure S2 for intermediate exploitation time). This has two consequences. First, the total number of offspring born in each patch is no longer the same as not all resources can be consumed in the time given. Instead, patches where individuals are better locally adapted can extract more resources and thus produce more offspring (lead-ing to hard selection; see Appendix C.2.4 for an analysis where exploitation time is short and selection is soft through an extra regulation step). Second, the absence of interference through resource consumption means that there cannot be any negative frequency-dependent selection within patches, i.e., no locally disruptive selection. In what follows, selection for polymorphism is therefore exclusively driven by spatially disruptive selection. To obtain analytical results, we assume that there are two types of patches showing a high degree of symmetry: specifically that the two types of patches occur at equal frequency (*π*_1_ *= π*_2_ *=* 1/2), that the distribution of resources within patches is Gaussian with the same variance 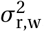, and the mean for patches of type 1 and 2 are given by 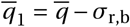 and 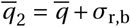, respectively, such that 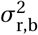 is the variance of patch means (Appendix C.2.1 for details on this model). This model can thus be seen as an extension to that of Meszéna et al. (1997) and of Ronce and Kirkpatrick (2001) who analyzed two-patch models under hard selection for local adaptation in the absence of within-patch resource differentiation (i.e. 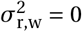 so that *E*_ST_ *=* 1) and of kin competition (infinite patch size *n* → ∞).

As with long exploitation time *T*, we find that the trait value 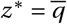 that matches the average resource is a singular strategy (Appendix C.2.2). A numerical analysis shows that provided between-patch resource differentiation *E*_ST_ is not too strong, *z*^*§*^ is an attractor of directional selection (Fig. 5a-c, solid colour regions). In other words, as long as patches are not too differentiated, the population initially converges to express 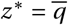. In this case, selection is disruptive and leads to polymorphism when

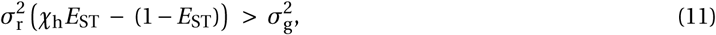

where *χ*_h_ ≥ 0 depends in a complicated manner on demographic parameters (on patch size *n*, dispersal probability *m*, and adult survival *γ*; eq. C54 in Appendix for details; Fig. 5a-c solid pink region for area where condition eq. 11 holds). Otherwise, selection is stabilising and the population remains fixed for 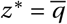 (Fig. 5a-c solid blue region for area where condition eq. 11 does not hold). The factor *χ*_h_ responds to demographic parameters in the same way as *χ*_1_ (eq. 10a), albeit for the fact that *χ*_h_ is larger by a small margin (Fig. S1). In the limit of large patch size and weak dispersal, the leading term of *χ*_h_ reduces to that of *χ*_1_ (shown in eq. 10a). In contrast to eq. (9), weaker resource differentiation *E*_ST_ always disfavors polymorphism in eq. (11) (Fig. 5a-c solid blue region). The reason for this is that here polymorphism is only driven by spatially disruptive selection. Variation of resources within patches weakens the strength of spatially disruptive selection because it allows individuals expressing the average resource trait 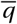 to exploit some resources in both patch types.

**Figure 5:**
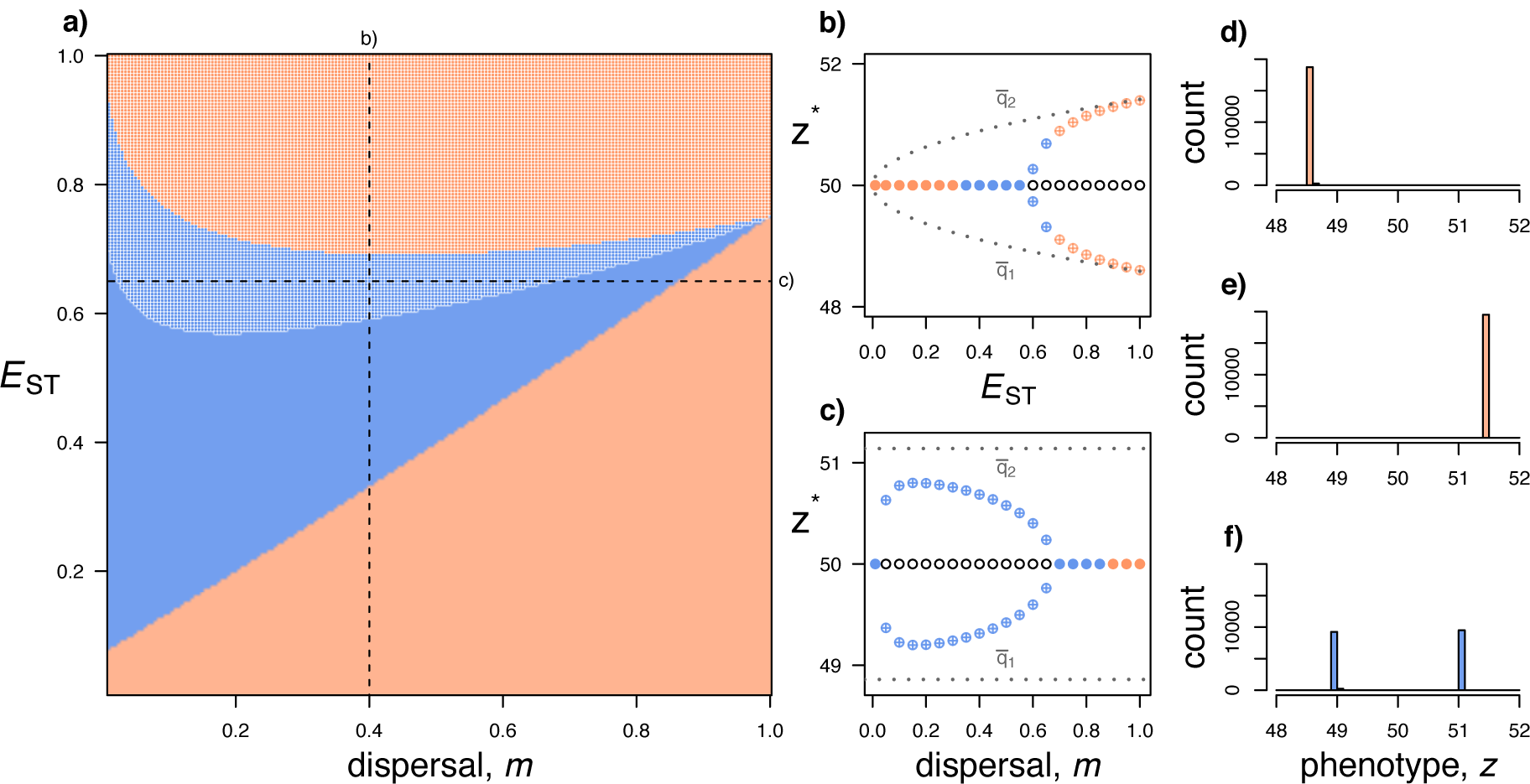
Evolutionary dynamics under hard selection. **a)** Combinations of resource differentiation *E*_ST_ and level of dispersal *m* that lead to stabilising (pink) or disruptive selection (blue) when exploitation time is short so that the number of offspring produced in each patch may vary (so that selection is hard, section 3.3 for details; other parameters: 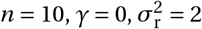, and 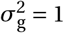; Appendix C.2 for analysis). Under stabilising selection, the population adapts to the average resource property 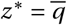 (solid pink) or to one of the two habitats depending on initial conditions (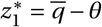 or 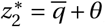, speckled pink). When selection is disruptive, the population first converges to 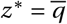 (solid blue) or to either 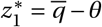 or 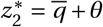 (speckled blue). Dashed lines indicate the values used in b) and c). **b)-c)** Bifurcation diagrams for the stability of singular strategies as a function of *E*_ST_ (in b with *m =* 0.65) and *m* (in c with *E*_ST_ *=* 0.4; other parameters: same as for a). White circles indicate singular strategies that are evolutionary repellors; pink circles indicate singular strategies that are attractors and for which selection is stabilising (solid: 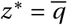; crossed: 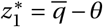 or 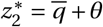); blue circles indicate singular strategies that are attractors and for which selection is disruptive (evolutionary branching points; solid: 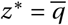; crossed: 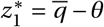 or 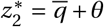). Dotted lines indicate the average resource property in each habitat, 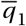 and 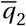. **d)-f)** Phenotypic distribution in simulated populations after 2,000 years of evolution when the population becomes adapted to a single habitat (when *E*_ST_ *=* 1 in d and e; with population initially monomorphic for *z =* 49 in d and *z =* 51 in e) or polymorphic (in f, where *E*_ST_ *=* 0.6). Other parameters: 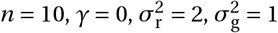 and *m =* 0.1.

Another contrasting result from the case where exploitation time *T* is long is that the singular strategy 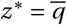 is no longer always an attractor of directional selection. In fact, when *E*_ST_ is sufficiently large, 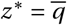 is an evolutionary repellor (Fig. 5a-b, speckled region) as long as dispersal is not extremely limited (Fig. 5c). When 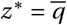 is a repellor, the population is attracted to either of two singular strategies: 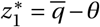 (for some *θ >* 0), which is closer to the average resource 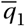 within patches of type 1; or 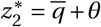, which is closer to the average resource 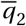 within patches of type 2. It is not possible to solve for 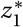 and 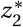 explicitly, but numerical explorations indicate that 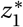 and 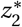 approach 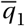 and 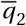, respectively, when *E*_ST_ increases and *m* is intermediate (Fig. 5b,c).

Depending on initial conditions, the population thus either converges to become more adapted to patches of type 1 (if the population initially expresses 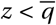) or to patches of type 2 (if the population initially expresses 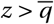). This is due to the fact that patches where individuals are better locally adapted send out more offspring. As a result, if individuals are initially fitter in one habitat, offspring with traits more adapted to these patches swamp the population, which then evolves to become more adapted to this habitat.

Investigating numerically the nature of selection when the population expresses 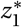 or 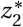 reveals that when patch differentiation *E*_ST_ is large, selection is stabilising at these singular points (Fig. 5a-c, speckled pink region), so that the population is “stuck” being adapted to a single habitat (Fig. 5d,e). When *E*_ST_ is intermediate, however, the population experiences disruptive selection and thus becomes polymorphic (Fig. 5a-c, speckled blue region). Individual-based simulations confirm this and show that eventually the population again consists of two morphs, each more adapted to one habitat (Fig. 5f). The endpoint of the dimorphic evolutionary dynamics does not depend on whether the population first converged to 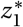 or 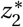. This shows that variation of resources within patches (so that *E*_ST_ *<* 1) allows the population to escape the evolutionary dead-end of being adapted to only one habitat. Similarly, Meszéna et al. (1997) found that intermediate patch divergence allowed the population to evolve polymorphism after first adapting to one patch type. Our model shows that these alternative routes to polymorphism can also be opened by variation of resources within patches.

## 4 Discussion

Our analyses indicate that polymorphism in a consumer trait emerges more readily when resources vary both within and between patches, in other words, when selection is simultaneously spatially and locally disruptive. Kin selection due to limited dispersal, by contrast, opposes polymorphism, especially when polymorphism is driven by locally disruptive selection. This is because local interactions among kin reduces the advantage of expressing alternative phenotypes to escape resource competition (Day, 2001; Ajar, 2003). Thus, although both spatially and locally disruptive selection contribute jointly to polymorphism, spatially disruptive selection is typically stronger because it is less hindered by kin competition and because of its interplay with dispersal. One broad conclusion is therefore that polymorphism results more easily from local adaptation than from local competition, although both contribute to the evolution of phenotypic variation.

Our finding that spatially and locally disruptive selection act in concert in promoting polymorphism contrasts with a previous suggestion that they may oppose one another (i.e. that polymorphism emerges less easily when both form of selection operate jointly, Day, 2000). This suggestion was drawn from a two-patch model of resource competition excluding kin selection (effectively assuming that patches are of infinite sizes). The phenomenological nature of Day (2000)’s model makes it difficult to pinpoint the biological reason for our contrasting results. While spatially and locally disruptive selection unfold from a microscopic ecological model in our study (in the spirit of Geritz and Kisdi 2004; our section 2.2), Day (2000) incorporates both types of selection independently by modifying dynamical equations of the Lotka-Volterra type. Locally disruptive selection is enforced by assuming that fecundity is limited by trait similarity with patch neighbours, and spatially disruptive selection by assuming that the carrying capacity of a patch decreases with the difference between a local phenotypic optima and the average trait expressed in that patch (eqs. 3-4 and 8 in Day, 2000). Although it is difficult to conceive an ecological scenario where trait expression has independent effects on fecundity and carrying capacity, Day (2000)’s results suggest that spatially and locally disruptive selection may not always work hand in hand towards polymorphism like in our model.

Whether selection leads to polymorphism is a different question from how selection shapes variation within and between patches once polymorphism has emerged. To this question, spatially and locally disruptive selection offer opposite answers: spatially disruptive selection favors local adaptation and phenotypic differentiation between patches, while locally disruptive selection favors variation within patches. Presumably, these antagonistic effects do not depend on the ecological scenario that leads to locally disruptive selection (which in our model is due to limiting similarity in resource consumption). For instance, Bolnick and Stutz (2017) analyses a two-patch population genetics model of a locus subject to: (i) negative frequency-dependent selection within patches whereby the rarer allele in a patch is favored in that patch to capture an ecological situation where parasites evolve to target the most abundant local type; and (ii) selection for local adaptation so that each allele is associated with greater fitness in a specific patch. Similarly to us, Bolnick and Stutz (2017) found that by increasing effective gene flow, negative frequency-dependent selection reduces allelic divergence among patches.

To quantify divergence among patches, we applied the popular *Q*_ST_ measure of trait differentiation. This measure has been widely used in comparison to neutral genetic differentiation 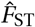 to infer the nature of selection in empirical studies (Leinonen et al., 2013). The commonly accepted interpretation of 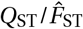 comparisons is that: 1) 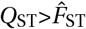 indicates spatially disruptive selection (i.e. local adaptation); 2) 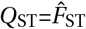 indicates neutral evolution; and 3) 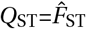 indicates spatially uniform selection (i.e. stabilising selection for the same trait value in all patches, e.g., Whitlock, 2008; Leinonen et al., 2013). In contrast, our results show that spatially uniform selection leads to 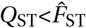. In line with this, a simulation study by Whitlock and Guillaume (2009) finds that spatially uniform selection leads at best to a weak signal of 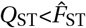 (when the phenotypic effects of mutations are large relative to selection), so that statistical methods have low detection power (their fig. 6 especially). Nevertheless, signals of 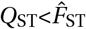 have been detected in several natural populations for a wide range of traits (e.g. Merilä and Crnokrak, 2001; McKay and Latta, 2002; Leinonen et al., 2008; Marin et al., 2020). Based on our results, an alternative suggestion is that these traits are in fact under locally disruptive selection, which causes greater divergence within patches than expected under random gene flow and thus 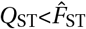 (as verbally argued by Lamy et al., 2012). In Snapdragon plants, for example, germination date shows lower levels of *Q*_ST_ relative to 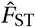 (Marin et al., 2020). Since germination date can mediate competition for resources (Elzinga et al., 2007), such pattern may in fact be due to locally disruptive selection. It would therefore be relevant to study more formally the statistical power of 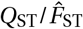 comparisons when frequency dependence promotes local polymorphism and thereby contributes to 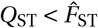.

**Figure 6:**
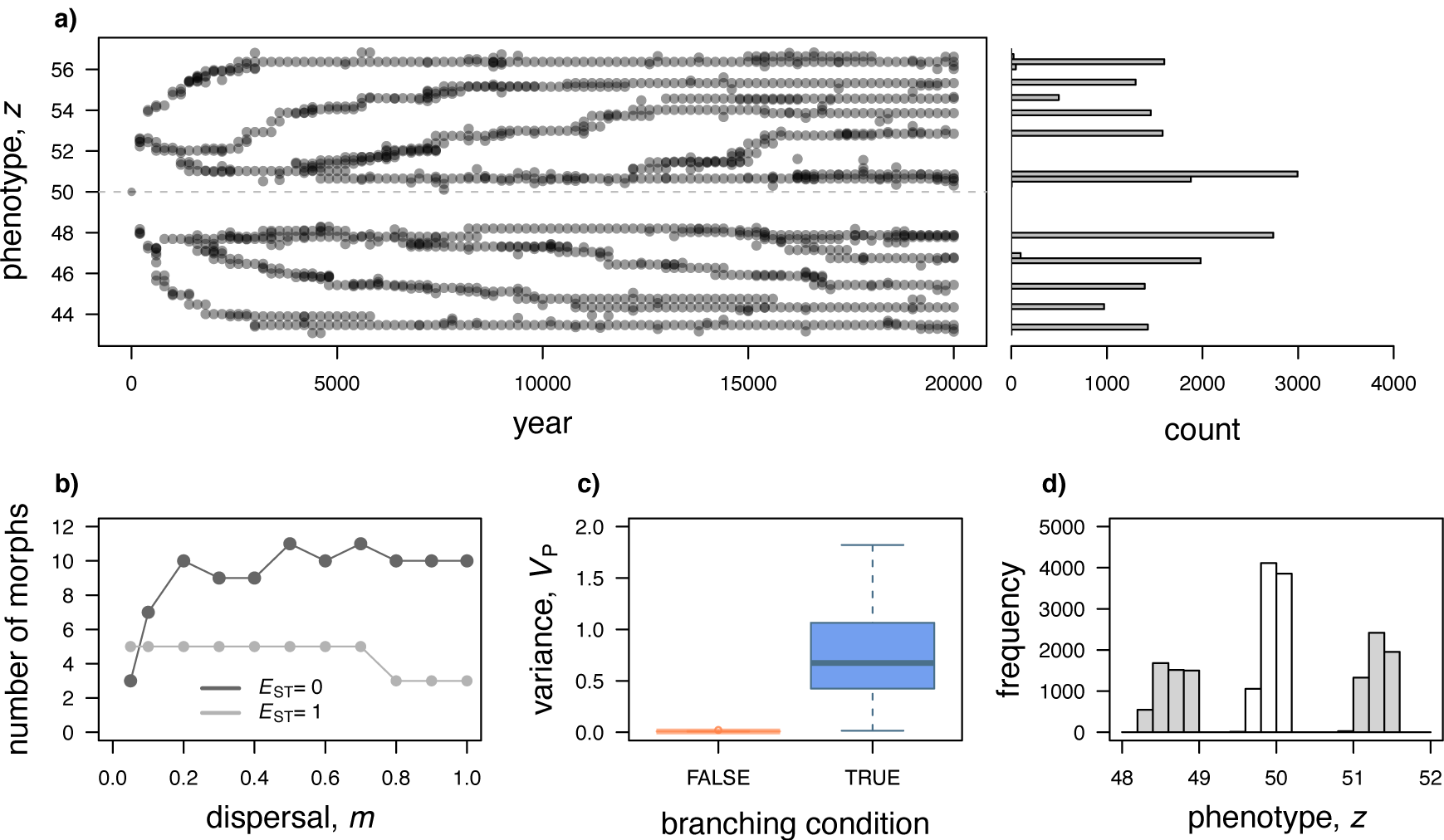
Emergence of multiple morphs and evolution under sexual reproduction. **a)** Simulated evolution of the consumer trait *z* when the resource distribution is wide relative to the degree of generalism, leading to the emergence of multiple morphs (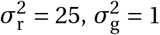; other parameters: *E*_ST_ *=* 0, *γ =* 0, *n =* 10, *m =* 0.5). **b)** The number of morphs after 20,000 years of simulated evolution as a function of dispersal for *E*_ST_ *=* 0 (in black) and *E*_ST_ *=* 1 (in gray; other parameters: 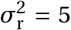 with 5 habitat types, 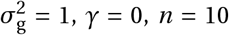). This shows that the number of morphs respectively increase and decrease with dispersal when selection is locally and spatially disruptive. **c)** Box-plots for the distributions of phenotypic variance *V*_P_ across multiple simulated populations of sexuals after 2,000 years of evolution when the branching condition eq. 9 is met (in blue) or not (in pink). Parameters varied: 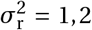; *E*_ST_ *=* 0, 0.5, 1; *m =* 0.05, 0.1, 0.2,…, 1.0; fixed parameters: 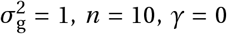. **d)** Phenotypic distribution in a simulated population of sexually reproducing individuals after 2,000 years of evolution (Parameters: 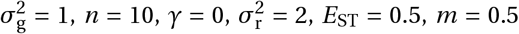). The most-diverged phenotypes are shown in gray while hybrids are shown in white.

Our simulations focus on situations where two morphs eventually coexist in the population, but there are cases in which more than two morphs may emerge and be maintained (Fig. 6a; Geritz et al., 1998). In the context of our model, multiple morphs may evolve when there are multiple patch types (i.e., the distribution in Fig. 1d top is multi-peaked) so that spatially disruptive selection favors local adaptation to many patch types; or when the degree of generalism 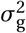 is much smaller than within-patch resource variation 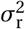, leading to strong locally disruptive selection to avoid competition for resources within patches. In principle, spatially disruptive selection favors as many morphs as there are patch types and locally disruptive selection as many to fill the local niches. But whether that many morphs actually evolve should depend on the degree of gene flow, with presumably fewer morphs under locally disruptive selection and more under spatially disruptive selection when dispersal is limited. Individual-based simulations confirm this, with the number of morphs increasing with dispersal when *E*_ST_ *=* 0 (Fig. 6b in black) and decreasing when *E*_ST_ *=* 1 (Fig. 6b in gray). To study this more definitively it would be necessary, though challenging, to perform an invasion analysis in subdivided polymorphic populations with finite patch size.

Our model assumes clonal reproduction but previous theory suggests that condition (9) for evolutionary branching and the emergence of highly differentiated morphs should not be affected by sexual reproduction (Kisdi and Geritz, 1999). To test this, we performed additional simulations where diploid individuals mate randomly within patches and the trait is determined by additive effects at one locus (Appendix D for details). These simulations show that trait variation is significantly greater when eq. (9) is satisfied than when it is not (Fig. 6c). In contrast to the case of clonal reproduction, the trait distribution in the population shows three instead of two morphs as mating among diverged genotypes creates intermediate hybrids (Fig. 6d). As these hybrids are less fit, selection should in turn favor mechanisms such as allelic dominance (Van Dooren, 1999) or assortative mating (Geritz and Kisdi, 2000) to prevent the formation of intermediate phenotypes (Slatkin, 1984; Kopp and Hermisson, 2006 for other mechanisms and Rueffler et al., 2006*a* for review). It would be interesting to extend our model to investigate the evolution of genetic and behavioural mechanisms that can avoid the production of unfit hybrids, especially because spatially and locally disruptive selection may favor the evolution of different mechanisms depending on dispersal and gene flow, with potential implications for speciation and diversity.

To conclude, our study reinforces the general notion that competition for resources is a major driver of adaptive diversification. It helps understand the conditions under which resource variation within and between habitats leads to consumer trait polymorphism, as well as how this polymorphism is distributed in the population. More broadly, by combining three fundamental sources of selection in spatially structured population, exploitation competition, local adaptation, and interactions among kin, our model is a step further towards integrating different strands of the theory of adaptation.

