## Appendix for "Foraging for locally and spatially varying resources: Where exploitation competition, local adaptation and kin selection meet"

#### A Fitness

Here, we specify how individuals collect resources and accumulate energy, and, in turn, how this determines individual fitness, which is the basis of our evolutionary analysis.

##### A.1 Collected resources and fecundity

To obtain the amount of energy accumulated by an individual before reproduction, we solve dynamical equations (4) for time $\tau$, and evaluate these solutions at the end of the period of exploitation (i.e., at $\tau = T$). We thus obtain that the amount of energy accumulated by a focal individual, say individual "1" among $n$ in a patch in state $s$, is

$$E_{1,s}(T) = E_s(z_1, \boldsymbol{z}_{-1}) = \beta \sum_{j=1}^{n_R} \left[ R_{j,s}(0) \left( 1 - e^{-T \sum_{k=1}^{n} \alpha(z_k, q_j)} \right) \right] \frac{\alpha(z_1, q_j)}{\sum_{k=1}^{n} \alpha(z_k, q_j)}, \tag{A1}$$

where recall $\beta$ is the conversion factor of resources into energy; $R_{j,s}(0) = R\pi_{j|s}$ is the initial amount of resource $j$ in a patch in state $s$ (given by eq. 5); and $\alpha(z_k, q_j)$ is the feeding rate of individual $k \in \{1, \ldots, n\}$ on resource $j$, which depends on the match between its phenotype $z_k$ and the quantitative property $q_j$ of the resource (eq. 3).

We assume that the fecundity of an individual increases linearly with the total amount of energy it has accumulated from resources (eq. 7), so that the fecundity of a focal individual 1 with trait $z_1$ when the other individuals of its patch have traits $\boldsymbol{z}_{-1} = (z_2, z_3, \dots, z_n)$ and its patch is in state $s$ is given by,

$$f_s(z_1, \boldsymbol{z}_{-1}) = kE_s(z_1, \boldsymbol{z}_{-1}), \tag{A2}$$

where $k$ is the conversion factor between energy and number of offspring produced.

#### A.2 Individual fitness

Given individual fecundity, we can now determine individual fitness, which is here defined as the expected number of successful offspring an individual produces over one full iteration of the life cycle (including itself if it survives). For our analysis specifically (which is based on Ohtsuki et al., 2020, see Appendix B below), we need to characterize the fitness function, $w_{s'|s}(z_1, \boldsymbol{z}_{-1}, z)$, which gives the expected number of offspring that settle in a patch in state $s'$ and that descend from an individual with trait $z_1$ in a patch in state $s$, where the patch neighbors to the focal express traits $\boldsymbol{z}_{-1} = (z_2, z_3, \dots, z_n)$ and the rest of the population is monomorphic for $z$.

##### A.2.1 Decomposing fitness

Following Ohtsuki et al. (2020), we first decompose individual fitness into two components,

$$w_{s'|s}(z_1, \boldsymbol{z}_{-1}, z) = \begin{cases} w^{\mathrm{p}}_{s|s}(z_1, \boldsymbol{z}_{-1}, z) + w^{\mathrm{d}}_{s|s}(z_1, \boldsymbol{z}_{-1}, z), & \text{for } s' = s, \\ w^{\mathrm{d}}_{s'|s}(z_1, \boldsymbol{z}_{-1}, z), & \text{for } s' \neq s, \end{cases} \tag{A3}$$

where $w^{\mathrm{d}}_{s'|s}(z_1, \boldsymbol{z}_{-1}, z)$ is the expected number of offspring that settles in non-natal patches of state $s'$, which we refer to as the dispersal component of fitness; and $w^{\mathrm{p}}_{s|s}(z_1, \boldsymbol{z}_{-1}, z)$ is the expected number of non-dispersing descendants (including the focal individual if it survives), which we refer to as the philopatric component. This component can be further decomposed into survival and reproduction components as

$$w^{\mathrm{p}}_{s|s}(z_1, \boldsymbol{z}_{-1}, z) = \gamma + n(1-\gamma)\phi^{\mathrm{p}}_{s|s}(z_1, \boldsymbol{z}_{-1}, z), \tag{A4}$$

where recall $\gamma$ is the probability of survival, and $\phi^{\mathrm{p}}_{s|s}(z_1, \boldsymbol{z}_{-1}, z)$ is the probability that an open breeding spot (of which there are $n(1-\gamma)$ on average before the regulation stage) is filled by a philopatric offspring of the focal individual (with $n\phi^{\mathrm{p}}_{s|s}(z_1, \boldsymbol{z}_{-1}, z)$ corresponding to $w^{pr}_{1,s|s}$ in Ohtsuki et al., 2020). Similarly, we write the dispersal component of fitness as,

$$w^{\mathrm{d}}_{s'|s}(z_1, \boldsymbol{z}_{-1}, z) = n(1-\gamma)\phi^{\mathrm{d}}_{s'|s}(z_1, \boldsymbol{z}_{-1}, z) \tag{A5}$$

where $\phi^{\mathrm{d}}_{s'|s}(z_1, \boldsymbol{z}_{-1}, z)$ is the probability that a dispersing offspring of the focal individual settles in an empty breeding spot in a patch of type $s'$ (note that $n\phi^{\mathrm{d}}_{s'|s}(z_1, \boldsymbol{z}_{-1}, z)$ corresponds to $w^{pr}_{1,s'|s}$ in Ohtsuki et al., 2020).

The probability that an offspring of the focal individual (with trait $z_1$) fills a spot in the focal patch is given by

$$\phi^{\mathrm{p}}_{s|s}(z_1, \boldsymbol{z}_{-1}, z) = \frac{(1-m) f_s(z_1, \boldsymbol{z}_{-1})}{(1-m)\sum_{i=1}^{n} f_s(z_i, \boldsymbol{z}_{-i}) + nmf(z)}, \tag{A6}$$

where

$$f(z) = \sum_{s\in\Omega} \pi_s f_s(z), \tag{A7}$$

is the average fecundity in a monomorphic population, which depends on

$$f_s(z) = \frac{k}{n}\sum_{j=1}^{n_{\mathrm{R}}} \beta R_{j,s}(0)\left(1 - e^{-Tn\alpha(z,q_j)}\right), \tag{A8}$$

the fecundity of an individual in a patch in state $s$ when all individuals in the patch express the same trait, $z$ (i.e., under neutrality), which is obtained from eq. (A2) by setting $z_1 = z_2 = \ldots = z_n = z$. Eq. (A6) thus consists of the ratio of the number of offspring of the focal that remain in their natal patch to the total number of offspring that enter competition, which is decomposed as the sum of those that are philopatric and those that come from other patches. Similarly, the probability that a dispersing offspring of the focal individual fills a spot in a patch in state $s'$ can be expressed as,

$$\phi^{\mathrm{d}}_{s'|s}(z_1, \boldsymbol{z}_{-1}, z) = \pi_{s'} \frac{m f_s(z_1, \boldsymbol{z}_{-1})}{n(1-m) f_{s'}(z) + nmf(z)}, \tag{A9}$$

where the numerator corresponds to the total number of offspring of the focal that disperse into patches in state $s'$, and the denominator to the total number of offspring that compete for a spot in such a patch.

##### A.2.2 Fitness components in a monomorphic population

As they will be relevant to our analysis, we also give here the components of fitness when the population is monomorphic for $z$. In this case, we obtain from eqs. (A6)-(A9) that

$$\phi^{\mathrm{p}}_{s|s}(z) = \frac{(1-m) f_s(z)}{(1-m) n f_s(z) + nmf(z)} \tag{A10}$$

for the philopatric component, and

$$\phi^{\mathrm{d}}_{s'|s}(z) = \pi_{s'} \frac{m f_s(z)}{(1-m) n f_{s'}(z) + nmf(z)} \tag{A11}$$

for the dispersal component.

### B Method

#### B.1 Evolution in two time scales

In this appendix, we describe the approach we take to model the evolution of the consumer trait $z$, assuming mutations are constantly occurring at a small rate and with small phenotypic effects. Under such assumptions, trait evolution can be decomposed into two timescales (Metz et al., 1996; Geritz et al., 1998; Dercole and Rinaldi, 2008). First, the trait evolves under directional selection, gradually evolving larger or smaller trait values under the influx of mutations. The population may thus attain a "convergence stable strategy", which is an attractor of directional evolution. Second, selection is either stabilising so that the population remains unimodaly distributed for the convergence stable strategy, or disruptive so that the trait becomes polymorphic and differentiated with individuals expressing either large or small traits. Our approach to understand directional and stabilizing/disruptive selection is based on an evolutionary invasion analysis for populations that are divided into non-homogeneous groups (Ohtsuki et al., 2020). We summarize this approach below with a focus on our model (see Ohtsuki et al., 2020 for more general problems, e.g. where the sizes of patches varies).

#### B.2 Directional selection

##### B.2.1 Selection gradient

The selection gradient, $S(z)$, tells us whether selection favors an increase (when $S(z) > 0$) or decrease (when $S(z) < 0$) in the trait when the population is monomorphic for $z$. Accordingly, a (locally) convergence stable strategy $z^*$ is such that

$$S(z^*) = 0, \tag{B1}$$

i.e., it is a "singular strategy" where directional selection vanishes, and

$$\left.\frac{dS(z)}{dz}\right|_{z=z^*} < 0, \tag{B2}$$

i.e., a population away from this singular strategy is attracted to it.

##### B.2.2 Selection gradient in patch-structured populations

It has been shown that for populations divided into heterogeneous patches that the selection gradient can be expressed in terms of individual fitness as,

$$S(z) = \sum_{s'\in\Omega}\sum_{s\in\Omega} v_{s'}(z)\left[\frac{\partial w_{s'|s,1}}{\partial z_1} + (n-1)r_{2,s}(z)\frac{\partial w_{s'|s,1}}{\partial z_2}\right] q_s(z) \tag{B3}$$

(e.g. eq. E.17 in Lehmann et al., 2016, eq. 32 in Ohtsuki et al., 2020), where we have used the short-hand notation

$$w_{s'|s,1} = w_{s'|s}(z_1, \boldsymbol{z}_{-1}, z) \tag{B4}$$

to denote the fitness of a focal individual, and here and hereafter all derivatives are evaluated in a monomorphic resident population at $z$ (i.e., where $z_i = z$ for all $i$). In eq. (B3), $q_s(z)$ is the probability that a randomly sampled individual from a resident lineage (i.e. whose members express $z$) resides in a patch in state $s$ (the frequency distribution among patch states in a resident population), which can be thought of as the contexts in which a mutation changing trait value can arise. The term within square brackets is the sum of the direct ($\partial w_{s'|s,1}/\partial z_1$) and relatedness-weighted indirect ($\partial w_{s'|s,1}/\partial z_2$) effects of varying trait expression on the expected number of offspring that an individual in a patch in state $s$ leaves in patches in state $s'$, where $r_{2,s}(z)$ is the probability that two randomly sampled individuals from a patch in state $s$ are identical-by-descent (pairwise relatedness) under neutrality. Hence, this term in square brackets can be thought of as the net effect on the expected number offspring that settle in a patch in state $s'$ descending from a parent in a patch in state $s'$, due to a change in the trait of all the members of the lineage of this parent. Finally, each offspring needs to be weighted by its reproductive value $v_{s'}(z)$, which is its asymptotic contribution to the future of the population in the absence of selection (and thus takes into account the demographic consequences of such offspring). The three quantities $q_s(z)$, $r_{2,s}(z)$, $v_{s'}(z)$ are thus evaluated under the assumption that the population is monomorphic for $z$. For a detailed derivation of eq. (B3), see Lehmann et al. (2016) or Ohtsuki et al. (2020).

Relatedness can be derived using standard identity-by arguments (e.g. Karlin and McGregor, 1968; Rousset, 2004). For our model, such an argument leads to a recursion which at equilibrium satisfies

$$r_{2,s}(z) = \gamma^2 r_{2,s}(z) + 2\gamma(1-\gamma)(1-d_s(z))r^{\mathrm{R}}_{2,s}(z) + (1-\gamma)^2(1-d_s(z))^2 r^{\mathrm{R}}_{2,s}(z), \tag{B5}$$

where

$$r^{\mathrm{R}}_{2,s}(z) = \frac{1}{n} + \frac{n-1}{n} r_{2,s}(z), \tag{B6}$$

is the probability that two individuals sampled with replacement (hence the superscript R) from a patch in state $s$ are identical-by-descent, and

$$d_s(z) = \frac{mf(z)}{(1-m)f_s(z) + mf(z)} \tag{B7}$$

is the backward probability of dispersal, i.e., the probability that an individual sampled in a patch of type $s$ is an immigrant in a population monomorphic for $z$. Solving eq. (B5) for $r_{2,s}(z)$ then gives

$$r_{2,s}(z) = \frac{2\gamma(1-d_s(z)) + (1-\gamma)(1-d_s(z))^2}{n(1+\gamma) - 2\gamma(n-1)(1-d_s(z)) - \left(1-\gamma\right)(n-1)(1-d_s(z))^2}, \tag{B8}$$

which agrees with eqs. (52) of Ohtsuki et al. (2020) that was derived with a different method. Meanwhile, the product between parental state distribution $q_s(z)$ and the offspring reproductive value $v_{s'}(z)$ appearing in eq. (B3) is given by,

$$v_{s'}(z)q_s(z) = \frac{\phi^{\mathrm{d}}_{s|s'}(z)}{(1-\gamma)\left(1-n\phi^{\mathrm{p}}_{s'|s'}(z)\right)\left(1-n\phi^{\mathrm{p}}_{s|s}(z)\right)} \Bigg/ \left(\sum_{s''\in\Omega} \frac{\phi^{\mathrm{d}}_{s''|s''}(z)}{(1-\gamma)\left(1-n\phi^{\mathrm{p}}_{s''|s''}(z)\right)^2}\right) \tag{B9}$$

(from eq. 40 in Ohtsuki et al., 2020).

##### B.2.3 Selection gradient with fecundity effects

Substituting for individual fitness $w_{s'|s}(z_1, \boldsymbol{z}_{-1}, z)$ in eq. (B3) with the explicit expression derived in Appendix A for our model (see also eq. I10 and eq. H10 in Ohtsuki et al., 2020 for more general results), we find after some re-arrangements that the selection gradient reduces to a single sum over patch states that can be written as

$$S(z) = \sum_{s\in\Omega} (1-\gamma)\left[F_s(z) - \Phi_s(z)^2\, r^{\mathrm{R}}_{2,s}(z)\, \overline{F}_s(z)\right] v_s(z)q_s(z), \tag{B10}$$

where $v_s(z)q_s(z)$ is the reproductive value of class $s$, i.e., the probability that an individual randomly sampled from the population descends from an individual reproducing in a patch in state $s$ (satisfying $\sum_{s\in\Omega} v_s(z)q_s(z) = 1$, e.g. Taylor, 1990; Rousset, 2004). Selection in patches of type $s$ then depends on the fraction of open breeding spots $(1-\gamma)$ and on

$$F_s(z) = \frac{\partial f_{s,1}}{\partial z_1} + (n-1)r_{2,s}(z)\frac{\partial f_{s,1}}{\partial z_2}, \tag{B11}$$

where we used

$$f_{s,1} = \frac{f_s(z_1, \boldsymbol{z}_{-1})}{f_s(z)}, \tag{B12}$$

for the fecundity of a focal individual relative to fecundity in a population monomorphic for $z$ (we drop the dependencies on phenotypes for the sake of brevity). Accordingly, $F_s(z)$ is the sum of the direct and relatedness-weighted fecundity effects of the trait in patches of type $s$ relative to $f_s(z)$. This indicates that selection tends to favor trait values that increase the fecundity of an individual and of its relatives.

But when a trait increases the fecundity of its bearer and/or of its relatives, this also leads to an increase in kin competition within patches which in turn tends to reduce the strength of selection. Such effect is captured in eq. (B10) by $\Phi_s(z)^2\, r^{\mathrm{R}}_{2,s}(z)\, \overline{F}_s(z)$ which consists of three terms. The first, $\Phi_s(z)^2$ gives the probability that two offspring born in the same patch of type $s$ compete with one another in a population monomorphic for $z$, since

$$\Phi_s(z) = 1 - d_s(z) \tag{B13}$$

is the probability that a randomly sampled individual in a patch in state $s$ is philopatric. Kin competition also increases with the probability $r^{\mathrm{R}}_{2,s}(z)$ that two offspring sampled before dispersal in a patch of type $s$ are identical-by-descent (given in eq. B6), and

$$\overline{F}_s(z) = \frac{\partial f_{s,1}}{\partial z_1} + (n-1)\frac{\partial f_{s,1}}{\partial z_2}, \tag{B14}$$

which is the total effect of the trait on relative fecundity, i.e., the effect on the fecundity of a focal individual in a patch of type $s$ if every individual in the patch increase their trait value infinitesimally. Altogether, eq. (B10) thus reflects the balance between the positive effects of a trait change when such a change increases fecundity and its negative indirect effects through increased kin competition.

It is useful to note that the class reproductive value can be written as

$$v_s(z)q_s(z) = K_s(z)\pi_s \tag{B15}$$

where

$$K_s(z) = \frac{f_s(z)}{f(z)} \frac{\frac{f_s(z)}{f(z)}(1-m)+m}{C^2_{\text{v,f}}(z)(1-m)+1} \tag{B16}$$

in which

$$C_{\text{v,f}}(z) = \sqrt{\sum_{s\in\Omega} \pi_s \left(\frac{f_s(z)}{f(z)} - 1\right)^2} \tag{B17}$$

is the coefficient of variation in fecundity among patch states, i.e., the ratio of the standard deviation to the mean fecundity. Hence, in the absence of fecundity variation across patches, $K_s(z) = 1$ for all $\in \Omega$.

#### B.3 Disruptive versus stabilizing selection

##### B.3.1 Disruptive selection coefficient

Once the population has converged to a singular strategy $z^*$ (so that satisfies eqs. B1-B2), whether selection is stabilising or disruptive depends on the sign of the coefficient of disruptive selection, which we denote generically as $H(z)$. Specifically, selection is stabilising and the population remains monomorphic for the convergence stable strategy $z^*$ when $H(z^*) < 0$, and conversely selection is disruptive leading to polymorphism when $H(z^*) > 0$ (Geritz et al., 1998; Dercole and Rinaldi, 2008; Rousset, 2004).

##### B.3.2 Disruptive selection coefficient in patch-structured populations

In a non-homogeneous group-structured population, the coefficient of disruptive selection has been shown to be composed of three biologically relevant terms,

$$H(z) = H_{\text{w}}(z) \;+\; H_{\text{q}}(z) \;+\; H_{\text{r}}(z) \tag{B18a}$$

(see eq. 22a and eq. 34a-c in Ohtsuki et al., 2020). The first,

$$\begin{aligned} H_{\mathrm{w}}(z) = &\frac{1}{2}\sum_{s\in\Omega}\sum_{s'\in\Omega} v_{s'}(z) \\ &\times\left[\frac{\partial^2 w_{s'|s,1}}{\partial z_1^2} + (n-1)r_{2,s}(z)\left\{\frac{\partial^2 w_{s'|s,1}}{\partial z_2^2} + 2\frac{\partial^2 w_{s'|s,1}}{\partial z_1 \partial z_2}\right\} + (n-1)(n-2)r_{3,s}(z)\frac{\partial^2 w_{s'|s,1}}{\partial z_2\partial z_3}\right] q_s(z), \end{aligned} \tag{B18b}$$

consists of direct and relatedness-weighted indirect second-order effects of the trait on fitness, where $r_{3,s}(z)$ is threeway relatedness: the probability that three individuals from the same patch in state $s$ are identical-by-descent in a population monomorphic for $z$; the second,

$$H_{\mathrm{q}}(z) = \sum_{s\in\Omega}\sum_{s'\in\Omega} v_{s'}(z)\left[\frac{\partial w_{s'|s,1}}{\partial z_1} + (n-1)r_{2,s}(z)\frac{\partial w_{s'|s,1}}{\partial z_2}\right] q_s^{(1)}(z), \tag{B18c}$$

depends on the first-order effect $q_s^{(1)}(z)$ of a trait change on the probability that a carrier of this change is in a patch of type $s$; and finally, the third,

$$H_{\mathrm{r}}(z) = \sum_{s\in\Omega}\sum_{s'\in\Omega} v_{s'}(z)\left[(n-1)r_{2,s}^{(1)}(z)\frac{\partial w_{s'|s,1}}{\partial z_2}\right] q_s(z), \tag{B18d}$$

captures first-order effects $r_{2,s}^{(1)}(z)$ of the trait on relatedness (i.e., the first-order effect of a trait change on the probability that a randomly sampled neighbour carrier of this change in a patch of type $s$ also expresses the trait change).

Threeway relatedness is found by solving the coalescent recursion,

$$r_{3,s}(z) = \gamma^3 r_{3,s}(z) + 3\gamma^2(1-\gamma)(1-d_s(z))r_{2:3,s}^{\mathrm{R}}(z) + (1-\gamma)^2(1-d_s(z))^2\left[3\gamma + (1-\gamma)(1-d_s(z))\right] r_{3,s}^{\mathrm{R}}(z), \tag{B19}$$

where

$$r_{2:3,s}^{\mathrm{R}}(z) = \frac{2}{n}r_{2,s}(z) + \frac{n-2}{n}r_{3,s}(z) \tag{B20}$$

is the probability that three individuals sampled randomly, two without replacement and one with replacement, from the same patch in state $s$ are identical-by-descent, and

$$r_{3,s}^{\mathrm{R}}(z) = \frac{1}{n^2} + 3\frac{1}{n}\frac{(n-1)}{n}r_{2,s}(z) + \frac{(n-1)}{n}\frac{(n-2)}{n}r_{3,s}(z), \tag{B21}$$

is the probability that three individuals sampled with replacement from the same patch in state $s$ are identical-by-descent. We do not present here the explicit solution of eq. (B19) even though it is straightforward to obtain, as the solution is complicated (see eqs. F32 of Ohtsuki et al., 2020 for this solution that was obtained by a different method but agrees with the present one). Eq. (B18) further depends on the trait effect on the distribution in patch states, which has been shown to be such that

$$
\begin{aligned}
v_{s'}(z)q_s^{(1)}(z) = \Bigg( & \frac{1}{(1/n)-\phi^{\mathrm{p}}_{s|s}(z)}\left[\frac{\partial \phi^{\mathrm{p}}_{s|s}(z_1,\boldsymbol{z}_{-1},z)}{\partial z_1} + (n-1)r_{2,s}(z)\frac{\partial \phi^{\mathrm{p}}_{s|s}(z_1,\boldsymbol{z}_{-1},z)}{\partial z_2}\right] - \\
& \sum_{s''\in\Omega}\frac{1}{(1/n)-\phi^{\mathrm{p}}_{s''|s''}(z)}\left[\frac{\partial \phi^{\mathrm{p}}_{s|s}(z_1,\boldsymbol{z}_{-1},z)}{\partial z_1} + (n-1)r_{2,s''}(z)\frac{\partial \phi^{\mathrm{p}}_{s''|s''}(z_1,\boldsymbol{z}_{-1},z)}{\partial z_2}\right]\Bigg)v_{s'}(z)q_s(z),
\end{aligned}
\tag{B22}
$$

(eq. 43 in Ohtsuki et al., 2020), as well as the trait effect on relatedness, which can be computed as,

$$
r^{(1)}_{2,s}(z) = \frac{2n^2 r_{2,s}(z)\left[\gamma+(1-\gamma)n\phi^{\mathrm{p}}_{s|s}(z)\right]}{2\gamma n\phi^{\mathrm{p}}_{s|s}(z)+(1-\gamma)\left(n\phi^{\mathrm{p}}_{s|s}(z)\right)^2}\left(r^{\mathrm{R}}_{2,s}(z)\frac{\partial \phi^{\mathrm{p}}_{s|s}(z_1,\boldsymbol{z}_{-1},z)}{\partial z_1} + (n-1)r^{\mathrm{R}}_{2:3,s}(z)\frac{\partial \phi^{\mathrm{p}}_{s|s}(z_1,\boldsymbol{z}_{-1},z)}{\partial z_2}\right)
\tag{B23}
$$

(eq. 44 in Ohtsuki et al., 2020).

##### B.3.3 Disruptive selection coefficient with fecundity effects

Plugging into eq. (B18) the expression for individual fitness $w_{s'|s}(z_1,\boldsymbol{z}_{-1},z)$ as well as its different components that we have derived for our model in Appendix A, we find after some re-arrangements that the coefficient of disruptive selection can be expressed as the average over patch states of three quantities that mirror the three terms of eq. (B18a),

$$
H(z) = \sum_{s\in\Omega}(1-\gamma)\left(H_{\mathrm{w},s}(z) + H_{\mathrm{q},s}(z) + H_{\mathrm{r},s}(z)\right)K_s(z)\pi_s
\tag{B24a}
$$

where $K_s(z)$ is given by eq. (B16). Those three quantities are,

$$
\begin{aligned}
H_{\mathrm{w},s}(z) =& \frac{1}{2}\Bigg[\left(\frac{\partial^2 f_{s,1}}{\partial z_1^2} + (n-1)\,r_{2,s}(z)\left(2\frac{\partial^2 f_{s,1}}{\partial z_1\partial z_2}+\frac{\partial^2 f_{s,1}}{\partial z_2^2}\right) + (n-1)\,(n-2)\,r_{3,s}(z)\frac{\partial^2 f_{s,1}}{\partial z_2\partial z_3}\right) \\
& - \Phi_s(z)^2\Bigg\{r^{\mathrm{R}}_{2,s}(z)\left(2\frac{\partial f_{s,1}}{\partial z_1}\overline{F}_s(z) + \frac{\partial^2 f_{s,1}}{\partial z_1^2} + (n-1)\,\frac{\partial^2 f_{s,1}}{\partial z_2^2}\right) \\
& + (n-1)r^{\mathrm{R}}_{2:3,s}(z)\left(2\frac{\partial f_{s,1}}{\partial z_2}\overline{F}_s(z) + 2\frac{\partial^2 f_{s,1}}{\partial z_1\partial z_2} + (n-2)\,\frac{\partial^2 f_{s,1}}{\partial z_2\partial z_3}\right)\Bigg\}\Bigg] + \Phi_s(z)^3 r^{\mathrm{R}}_{3,s}(z)\overline{F}_s(z)^2 \\
H_{\mathrm{q},s}(z) =& \frac{1-d_s(z)}{d_s(z)}\left(F_s(z) - r^{\mathrm{R}}_{2,s}(z)\,\Phi_s(z)\,\overline{F}_s(z)\right)\left(F_s(z) - r^{\mathrm{R}}_{2,s}(z)\,\Phi_s(z)^2\,\overline{F}_s(z)\right) \\
H_{\mathrm{r},s}(z) =& (n-1)\,r^{(1)}_{2,s}(z)\left(\frac{\partial f_{s,1}}{\partial z_2} - \frac{1}{n}\Phi_s(z)^2\,\overline{F}_s(z)\right),
\end{aligned}
\tag{B24b}
$$

where we have used notations introduced in sections B.2 and B.3.2 (see eqs. I13-I17 and H13-H17 in Ohtsuki et al., 2020 for more general results on the coefficient of disruptive selection on traits with fecundity effects). Further, substituting for the philopatric component of fitness (eqs. A6, A10) into (B23), we obtain that the trait effect on relatedness can be expressed in terms of fecundity effects as

$$
r^{(1)}_{2,s}(z) = 2n\; r_{2,s}(z)\frac{\gamma+(1-\gamma)\,(1-d_s(z))}{2\gamma+(1-\gamma)\,(1-d_s(z))}\left[r^{\mathrm{R}}_{2,s}(z)\frac{\partial f_{s,1}}{\partial z_1} + (n-1)r^{\mathrm{R}}_{2:3,s}(z)\frac{\partial f_{s,1}}{\partial z_2} - r^{\mathrm{R}}_{3,s}(z)\Phi_s(z)\overline{F}_s(z)\right].
\tag{B25}
$$

### C Analyses

In this appendix, we use the framework described in appendix B to investigate the evolution of the consumer trait $z$ and derive our results presented in the main text.

#### C.1 Baseline scenario: long exploitation time

As a baseline, we assume that the time for consumption $T$ within generations is long, or more specifically, long enough so that individuals have time to consume all available resources. This assumption allows our mathematical analysis to go further.

##### C.1.1 Fecundity

Letting $T \to \infty$ in eqs. (A1)–(A2), we obtain that the fecundity of the focal individual simplifies to

$$f_s(z_1, \boldsymbol{z}_{-1}) = k\beta R \sum_{j=1}^{n_\mathrm{R}} \pi_{j|s} \frac{\alpha(z_1, q_j)}{\sum_{k=1}^{n} \alpha(z_k, q_j)}. \tag{C1}$$

In this case, the sum of the fecundities of all individuals in the patch reads as

$$\sum_{i=1}^{n} f_s(z_i, \boldsymbol{z}_{-i}) = k\beta R \sum_{j=1}^{n_\mathrm{R}} \pi_{j|s} \sum_{i=1}^{n} \frac{\alpha(z_i, q_j)}{\sum_{k=1}^{n} \alpha(z_i, q_j)} = k\beta R = f_{\max}, \tag{C2}$$

i.e., each patch produces the same number of offspring $f_{\max} = k\beta R$ (which is the maximum possible fecundity for an individual in the presence of trait variation). Selection therefore is always soft in this model (the case of hard selection is explored later).

In population monomorphic for $z$ (where $z_i = z$), the fecundity of an individual in a patch in state $s$ is then simply,

$$f_s(z) = \frac{f_{\max}}{n}, \tag{C3}$$

which is thus also equal to the average fecundity in the population,

$$f(z) = \sum_{s \in \Omega} \pi_s f_s(z) = \frac{f_{\max}}{n}. \tag{C4}$$

As a result, the coefficient of variation in fecundity across patch types vanishes:

$$C_{\mathrm{v,f}}(z) = \sqrt{\sum_{s \in \Omega} \pi_s \left( \frac{f_s(z)}{f(z)} - 1 \right)^2} = 0. \tag{C5}$$

This further entails that the quantity $K_s(z)$ (eq. B16), which is relevant to both selection coefficients (eqs. B10 and B18) reduces to

$$K_s(z) = 1. \tag{C6}$$

Relative fecundity of a focal individual, meanwhile, is obtained by substituting eq. (C1) and eq. (C3) into eq. (B12) to get

$$f_{s,1} = \frac{f_s(z_1, \mathbf{z}_{-1})}{f_s(z)} = \sum_{j=1}^{n_R} \pi_{j|s} \frac{\alpha(z_1, q_j)}{\sum_{k=1}^{n} \alpha(z_k, q_j)/n}. \tag{C7}$$

We can then use the above quantities to investigate how selection shapes the consumer trait $z$ according to the framework described in appendix B.

##### C.1.2 Directional selection

Substituting eq. (3), namely, $\alpha(z_i, q_j) = \exp\left(-(z_i - q_j)^2/(2\sigma_g^2)\right)$, into eq. (C7), we find that the direct (relative) fecundity effect is

$$\frac{\partial f_{s,1}}{\partial z_1} = -\sum_{j=1}^{n_R} \pi_{j|s} \frac{n-1}{n} \frac{z - q_j}{\sigma_g^2} = -\frac{n-1}{n} \frac{z - \overline{q}_s}{\sigma_g^2}, \tag{C8}$$

where recall $\overline{q}_s = \sum_{j=1}^{n_R} \pi_{j|s} q_j$ is the average resource property in a patch of type $s$. Similarly, the indirect fecundity effect reads as

$$\frac{\partial f_{s,1}}{\partial z_2} = \frac{1}{n} \frac{z - \overline{q}_s}{\sigma_g^2}. \tag{C9}$$

Accordingly, the sum of the direct and relatedness-weighted indirect fecundity effects (eq. B11) is

$$F_s(z) = -\frac{n-1}{n} \frac{z - \overline{q}_s}{\sigma_g^2} \left[1 - r_2\right], \tag{C10}$$

where pairwise relatedness $r_2$ here is independent from the state of the patch state $s$ and the evolving trait $z$, i.e.,

$$r_2 = r_{2,z}(z) = \frac{2\gamma(1-m) + (1-\gamma)(1-m)^2}{n(1+\gamma) - 2\gamma(n-1)(1-m) - \left(1-\gamma\right)(n-1)(1-m)^2} \tag{C11}$$

for all patch state $s$ (to see this, substitute eqs. C3 and C4 into eq. B7 which is in turn substituted into eq. B8). The total fecundity effect (eq. B14) meanwhile vanishes,

$$\overline{F}_s(z) = 0. \tag{C12}$$

Substituting eqs. (C10) and (C12) into the selection gradient eq. (B10) and using eq. (C6), we obtain

$$S(z) = -\sum_{s \in \Omega} (1-\gamma)\pi_s \frac{n-1}{n} \frac{z - \overline{q}_s}{\sigma_g^2} \left[1 - r_2\right] = -\left[1 - r_2\right](1-\gamma) \frac{n-1}{n} \frac{z - \overline{q}}{\sigma_g^2}, \tag{C13}$$

where recall $\overline{q} = \sum_{s \in \Omega} \pi_s \overline{q}_s$ is the the global average resource property. It is immediate from eq. (C13) that the unique singular strategy $z^*$ (eq. B1) is

$$z^* = \overline{q} \tag{C14}$$

and that this strategy is convergence stable (eq. B2) since

$$\left.\frac{dS(z)}{dz}\right|_{z=z^*} = -\left[1-r_2\right](1-\gamma)\frac{n-1}{n}\frac{1}{\sigma_{\mathrm{g}}^2} < 0. \tag{C15}$$

##### C.1.3 Disruptive selection

We now compute the disruptive selection coefficient from eq. (B24). To that end, consider first the direct second-order fecundity effect at the singular strategy $z^* = \overline{q}$:

$$\frac{\partial^2 f_{s,1}}{\partial z_1^2} = \sum_{j=1}^{n_{\mathrm{R}}} \pi_{j|s}\frac{n-1}{n}\frac{1}{\sigma_{\mathrm{g}}^2}\left(\frac{n-2}{n}\frac{\left(q_j-\overline{q}\right)^2}{\sigma_{\mathrm{g}}^2}-1\right), \tag{C16}$$

which averaged over environmental states as in eq. (B24), becomes

$$\sum_{s\in\Omega}(1-\gamma)K_s(z)\frac{\partial^2 f_{s,1}}{\partial z_1^2}\pi_s = (1-\gamma)\frac{n-1}{n}\frac{1}{\sigma_{\mathrm{g}}^2}\left(\frac{n-2}{n}\frac{\sigma_{\mathrm{r,w}}^2+\sigma_{\mathrm{r,b}}^2}{\sigma_{\mathrm{g}}^2}-1\right), \tag{C17}$$

where we used $K_s(z) = 1$ (eq. (C6)) and the fact that $\sum_{s\in\Omega}\sum_{j=1}^{n_{\mathrm{R}}}\pi_{j|s}\pi_s(q_j-\overline{q})^2 = \sigma_{\mathrm{r,w}}^2+\sigma_{\mathrm{r,b}}^2 = \sigma_{\mathrm{r}}^2$ is the total variance in resource property. Following the same procedure as the one used for eqs. (C16)-(C17), we readily obtain the remaining relevant second-order fecundity effects for the disruptive selection coefficient (eq. B24):

$$\begin{aligned}
\sum_{s\in\Omega}(1-\gamma)K_s(z)\frac{\partial^2 f_{s,1}}{\partial z_2^2}\pi_s &= (1-\gamma)\frac{1}{n}\frac{1}{\sigma_{\mathrm{g}}^2}\left(1-\frac{n-2}{n}\frac{\sigma_{\mathrm{r,w}}^2+\sigma_{\mathrm{r,b}}^2}{\sigma_{\mathrm{g}}^2}\right)\\
\sum_{s\in\Omega}(1-\gamma)K_s(z)\frac{\partial^2 f_{s,1}}{\partial z_1\partial z_2}\pi_s &= -(1-\gamma)\frac{1}{n}\frac{1}{\sigma_{\mathrm{g}}^2}\frac{n-2}{n}\frac{\sigma_{\mathrm{r,w}}^2+\sigma_{\mathrm{r,b}}^2}{\sigma_{\mathrm{g}}^2}\\
\sum_{s\in\Omega}(1-\gamma)K_s(z)\frac{\partial^2 f_{s,1}}{\partial z_2\partial z_3}\pi_s &= (1-\gamma)\frac{2}{n}\frac{1}{\sigma_{\mathrm{g}}^2}\frac{1}{n}\frac{\sigma_{\mathrm{r,w}}^2+\sigma_{\mathrm{r,b}}^2}{\sigma_{\mathrm{g}}^2}.
\end{aligned} \tag{C18}$$

Using eqs. (C17)-(C18), we find that the two total effects,

$$\begin{aligned}
\sum_{s\in\Omega}(1-\gamma)K_s(z)\left(\frac{\partial^2 f_{s,1}}{\partial z_1^2}+(n-1)\frac{\partial^2 f_{s,1}}{\partial z_2^2}\right)\pi_s = 0,\\
\sum_{s\in\Omega}(1-\gamma)K_s(z)\left(2\frac{\partial^2 f_{s,1}}{\partial z_1 z_2}+(n-2)\frac{\partial^2 f_{s,1}}{\partial z_2 z_3}\right)\pi_s = 0,
\end{aligned} \tag{C19}$$

reduce to zero. As a result (and using eqs. (C10)-(C12)), the first term $H_{\mathrm{w}}(z) = \sum_{s\in\Omega}(1-\gamma)K_s(z)H_{\mathrm{w},s}(z)\pi_s$ of the disruptive selection coefficient reduces to

$$H_{\mathrm{w}}(z) = (1-\gamma)\frac{n-1}{n}\frac{1}{\sigma_{\mathrm{g}}^2}\frac{1}{2}\left(\frac{n-2}{n}(1-L)\frac{\sigma_{\mathrm{r,b}}^2+\sigma_{\mathrm{r,w}}^2}{\sigma_{\mathrm{g}}^2}-(1-r_2)\right), \tag{C20}$$

where

$$L = 3r_2 - 2r_3 \tag{C21}$$

is the probability that at least two out of three individuals are identical-by-descent (so that $1-L$ is the probability that none out of three are identical-by-descent ), with

$$r_3 = r_{3,s}(z) \quad \text{for all } s, \tag{C22}$$

as the probability of identity-by-descent of three individuals from the same patch. Like pairwise relatedness (eq. C11), $r_3$ is independent from patch state and the evolving trait here. It is found by solving

$$r_3 = \gamma^3 r_3 + 3\gamma^2(1-\gamma)(1-m)r_{2:3}^{\mathrm{R}} + (1-\gamma)^2(1-m)^2\left[3\gamma + (1-\gamma)(1-m)\right] r_3^{\mathrm{R}}, \tag{C23}$$

where

$$\begin{aligned} r_{2:3}^{\mathrm{R}} &= \frac{2}{n} r_2 + \frac{n-2}{n} r_3 \\ r_3^{\mathrm{R}} &= \frac{1}{n^2} + 3\frac{1}{n}\frac{(n-1)}{n} r_2 + \frac{(n-1)}{n}\frac{(n-2)}{n} r_3 \end{aligned} \tag{C24}$$

(obtained by substituting eqs. C3 and C4 into eq. B7 which is in turn substituted into eq. B19).

Plugging eqs. (B7), (B25), and eqs. (C9)-(C12) into eq. (B24), we find after some re-arrangements that the other two terms of the disruptive selection coefficient, $H_{\mathrm{q}}(z) = \sum_{s\in\Omega}(1-\gamma)K_s(z)H_{\mathrm{q},s}(z)\pi_s$ and $H_{\mathrm{r}}(z) = \sum_{s\in\Omega}(1-\gamma)K_s(z)H_{\mathrm{r},s}(z)\pi_s$, read as,

$$H_{\mathrm{q}}(z) = (1-\gamma)\frac{(n-1)}{n}\frac{1}{\sigma_{\mathrm{g}}^2}\left(\frac{(n-1)}{n}\frac{(1-m)}{m}(1-r_2)^2\frac{\sigma_{\mathrm{r,b}}^2}{\sigma_{\mathrm{g}}^2}\right) \tag{C25}$$

and

$$H_{\mathrm{r}}(z) = -(1-\gamma)\frac{(n-1)}{n}\frac{1}{\sigma_{\mathrm{g}}^2}\left(2(n-1)\frac{(1-(1-\gamma)m)}{(1-(1-\gamma)m+\gamma)} r_2\left(\frac{1-L}{n} + r_2 - r_3\right)\frac{\sigma_{\mathrm{r,b}}^2}{\sigma_{\mathrm{g}}^2}\right). \tag{C26}$$

Summing eqs. (C20), (C25), and (C26), we eventually obtain that the coefficient of disruptive selection at $z = \overline{q}$ can be expressed as

$$H(z) = \frac{1-\gamma}{\sigma_{\mathrm{g}}^2}\frac{n-1}{n}\left(\chi_a\frac{\sigma_{\mathrm{r,b}}^2}{\sigma_{\mathrm{g}}^2} + \chi_b\frac{\sigma_{\mathrm{r,w}}^2}{\sigma_{\mathrm{g}}^2} - \chi_c\right), \tag{C27}$$

where

$$\begin{aligned} \chi_a &= \frac{n-1}{n}\frac{1-m}{m}(1-r_2)^2 - 2(n-1)\frac{1-(1-\gamma)m}{1-(1-\gamma)m+\gamma}\left(\frac{1-L}{n} + r_2 - r_3\right) r_2 + \chi_b \\ \chi_b &= \frac{n-2}{n}\frac{1}{2}(1-L) \\ \chi_c &= \frac{1-r_2}{2}. \end{aligned} \tag{C28}$$

Substituting eq. (C27) into the branching condition, $H(z) > 0$, we obtain after some algebra condition (9) of the main text, namely

$$\frac{\sigma_{\mathrm{r}}^2}{\sigma_{\mathrm{g}}^2}\left(\chi_1 E_{\mathrm{ST}} + \chi_2(1-E_{\mathrm{ST}})\right) > 1, \tag{C29}$$

where

$$\chi_1 = \frac{\chi_a}{\chi_c} \quad \text{and} \quad \chi_2 = \frac{\chi_b}{\chi_c}. \tag{C30}$$

After inserting the expressions for the relatedness coefficients into eq. (C28) we obtain from eq. (C30) the definitions given by eq. (10) in the main text for $\chi_1$ and $\chi_2$ in the limit where patches are large and dispersal is weak (i.e., where $n \to \infty$ and $m \to 0$ such that $nm$ remains constant). Note that when dispersal is complete ($m = 1$) so individuals do not compete with related individuals ($r_2 = 0$ and $r_3 = 0$), $\chi_1$ and $\chi_2$ in eq. (C30) both reduce to

$$\chi_1 = \chi_2 = 1 - \frac{2}{n}, \tag{C31}$$

which increases to one as patch size increases.

#### C.2 Short exploitation time

Here we analyse our model when exploitation time $T$ is short and derive the results described in section 3.3 of the main text.

##### C.2.1 Fecundity

To obtain the fecundity of a focal individual expressing trait $z_1$ in a patch of type $s$, we first Taylor expand eq. (A1) around $T = 0$:

$$E_s(z_1, \boldsymbol{z}_{-1}) = \beta \sum_j R_{j,s}(0) T \alpha(z_1, q_j) + \mathcal{O}(T^2). \tag{C32}$$

Note how this depends only on the trait $z_1$ expressed by the focal individual (i.e., it is independent from the traits of other patch members). This is because exploitation time is too short for competition to affect the focal's resource uptake. Nonetheless, there is still competition for breeding spots whose outcome depends on the amount of resources collected.

Because the general case where resource distribution is arbitrary (given by $\pi_{j|s}$ and $\pi_s$), is too complicated, we assume that resources within patches follow a Normal distribution. In other words, we assume that there is an effectively infinite number of resource types that are continuously distributed such that the quantity of a resource $j$ with property $q_j$ in a patch of type $s$ is,

$$R_{j,s}(0) = R \frac{1}{\sqrt{2\pi\sigma^2_{\text{r,w}}}} \exp\left(-\frac{1}{2}\frac{(q_j - \overline{q}_s)^2}{\sigma^2_{\text{r,w}}}\right), \tag{C33}$$

where recall $\overline{q}_s$ is the average resource property in a patch of type $s$ (eq. C33 is the continuous analogue to eq. 5 of the main text). The fecundity of a focal individual is then calculated by plugging eq. (C33) into eq. (C32) which in turn is integrated over all resource types (rather than summed as in eq. 7 since the resource distribution is continuous), i.e.

$$f_s(z_1, \boldsymbol{z}_{-1}) = f_{\max} \int_{-\infty}^{\infty} \frac{1}{\sqrt{2\pi\sigma_{\mathrm{r,w}}^2}} \exp\left(-\frac{1}{2}\frac{(q_j - \overline{q}_s)^2}{\sigma_{\mathrm{r,w}}^2}\right) \alpha(z_1, q_j)\, \mathrm{d}q_j = f_{\max} \frac{\sqrt{\sigma_{\mathrm{g}}^2}}{\sqrt{\sigma_{\mathrm{g}}^2 + \sigma_{\mathrm{r,w}}^2}} \exp\left(-\frac{\left(z_1 - \overline{q}_s\right)^2}{2\left(\sigma_{\mathrm{g}}^2 + \sigma_{\mathrm{r,w}}^2\right)}\right), \tag{C34}$$

where $f_{\max} = k\beta RT$ is the maximum fecundity in this model and the function $\alpha(z_1, q_j)$ is given by eq. (3). From eq. (C34), fecundity in patch of type $s$ when the population is monomorphic for $z$ is then,

$$f_s(z) = f_{\max} \frac{\sqrt{\sigma_{\mathrm{g}}^2}}{\sqrt{\sigma_{\mathrm{g}}^2 + \sigma_{\mathrm{r,w}}^2}} \exp\left(-\frac{\left(z - \overline{q}_s\right)^2}{2\left(\sigma_{\mathrm{g}}^2 + \sigma_{\mathrm{r,w}}^2\right)}\right), \tag{C35}$$

so that we obtain

$$f_{s,1} = \frac{f_s(z_1, \boldsymbol{z}_{-1})}{f_s(z)} = \exp\left(-\frac{\left(z_1 - \overline{q}_s\right)^2 - \left(z - \overline{q}_s\right)^2}{2\left(\sigma_{\mathrm{g}}^2 + \sigma_{\mathrm{r,w}}^2\right)}\right), \tag{C36}$$

for the relative fecundity of the focal individual.

To derive further relevant quantities to our analysis, we make use of our assumption given in section 3.3 that there are two patch types, 1 and 2. These occur in equal frequency ($\pi_1 = \pi_2 = 1/2$) and differ in the average property of the resources they hold according to

$$\begin{aligned} \overline{q}_1 &= \overline{q} - \sigma_{\mathrm{r,b}} \\ \overline{q}_2 &= \overline{q} + \sigma_{\mathrm{r,b}}, \end{aligned} \tag{C37}$$

where $\overline{q}$ is the global average of resource property so that $\sigma_{\mathrm{r,b}}^2 = \left(\overline{q}_1 - \overline{q}\right)^2/2 + \left(\overline{q}_2 - \overline{q}\right)^2/2$ is the between patch variance in resource property, as required. In this case, the average fecundity in a monomorphic population reads as,

$$f(z) = f_{\max} \frac{1}{2} \frac{\sqrt{\sigma_{\mathrm{g}}^2}}{\sqrt{\sigma_{\mathrm{g}}^2 + \sigma_{\mathrm{r,w}}^2}} \left[\exp\left(-\frac{\left(z - \overline{q} - \sigma_{\mathrm{r,b}}\right)^2}{2\left(\sigma_{\mathrm{g}}^2 + \sigma_{\mathrm{r,w}}^2\right)}\right) + \exp\left(-\frac{\left(z - \overline{q} + \sigma_{\mathrm{r,b}}\right)^2}{2\left(\sigma_{\mathrm{g}}^2 + \sigma_{\mathrm{r,w}}^2\right)}\right)\right]. \tag{C38}$$

Plugging eqs. (C35) and (C38) into eq. (B17), we obtain

$$C_{\mathrm{v,f}}^2(z) = \tanh\left(-\frac{(z - \overline{q})\sigma_{\mathrm{r,b}}}{\sigma_{\mathrm{g}}^2 + \sigma_{\mathrm{r,w}}^2}\right)^2 \tag{C39}$$

for the coefficient of variation in fecundity in a monomorphic population, where tanh is the hyperbolic tangent function.

##### C.2.2 Directional selection

To compute the selection gradient $S(z)$ (from eq. B10), we first obtain from eq. (C36) the direct (relative) fecundity effect,

$$\frac{\partial f_{s,1}}{\partial z_1} = -\frac{z-\overline{q}_s}{\sigma_{\mathrm{g}}^2+\sigma_{\mathrm{r,w}}^2}, \tag{C40}$$

as well as the indirect fecundity effect,

$$\frac{\partial f_{s,1}}{\partial z_2} = 0. \tag{C41}$$

Accordingly, the two relevant total effects $F_s(z)$ (eq. B11) and $\overline{F}_s(z)$ (eq. B14) that appear in the selection gradient reduce to

$$F_s(z) = \overline{F}_s(z) = \frac{\partial f_{s,1}}{\partial z_1}. \tag{C42}$$

The selection gradient (eq. B10) thus simplifies to

$$S(z) = \frac{1-\gamma}{2}\sum_{s=1}^{2} K_s(z)\left(-\frac{z-\overline{q}_s}{\sigma_{\mathrm{g}}^2+\sigma_{\mathrm{r,w}}^2}\right)\left(1-\Phi_s(z)^2\, r_{2,s}^{\mathrm{R}}(z)\right), \tag{C43}$$

where we have used our assumption that there are two path types occurring at equal frequency (i.e., $\pi_1 = \pi_2 = 1/2$). The mathematical expressions for $K_s(z)$, $\Phi_s(z)$, and $r_{2,s}^{\mathrm{R}}(z)$ in eq. (C43) are then found using the different fecundity functions derived in section C.2.1 (plugged into eqs. B6-B8, B16 and B13). Although straightforward, this operation leads to an unsightly equation that is too complex to give much intuition on directional selection. We therefore do not present this equation and rather analyse the selection gradient numerically. This generates our results shown in Fig. 5 (i.e., we find singular strategies $z^*$ and test whether these are convergence stable using eqs. B1 and B2 for various parameter combinations shown in Fig. 5).

To show that $z^* = \overline{q}$ is a singular strategy, consider first that when the population is monomorphic for $z = \overline{q}$, one has $K_s(z) = 1$, as well as,

$$1-\Phi_s(z)^2\, r_{2,s}^{\mathrm{R}}(z) = 1-(1-m)^2\left(\frac{1}{n}+\frac{n-1}{n}r_2\right) \tag{C44}$$

with $r_2$ from eq. (C11) (when $z = \overline{q}$). The expression for the selection gradient (eq. C43) then reduces to

$$S(z) = -(1-\gamma)\left(1-(1-m)^2\left(\frac{1}{n}+\frac{n-1}{n}r_2\right)\right)\frac{(z-\overline{q})}{\sigma_{\mathrm{g}}^2+\sigma_{\mathrm{r,w}}^2}, \tag{C45}$$

indicating that $z^* = \overline{q}$ is always a singular strategy (where $S(z) = 0$). Our numerical analysis of the selection gradient, however, reveals that $z^* = \overline{q}$ is not always convergence stable. When it is convergence stable, it is the only singular strategy. But when $z^* = \overline{q}$ is not convergence stable, two additional singular strategies exist, one close to $\overline{q}_1$ and another close to $\overline{q}_2$ (Fig. 5). In this case convergence to either singularity depends on initial condition.

##### C.2.3 Disruptive selection

We now compute the disruptive selection coefficient (eq. B24). To do so, we first obtain from eq. (C36) the direct second-order effect,

$$\frac{\partial^2 f_{s,1}}{\partial z_1^2} = \left(\frac{z-\overline{q}_s}{\sigma_{\rm g}^2+\sigma_{\rm r,w}^2}\right)^2 - \frac{1}{\sigma_{\rm g}^2+\sigma_{\rm r,w}^2}. \tag{C46}$$

From eq. (C36) still, we further see that all indirect second-order effects reduce to zero,

$$\frac{\partial^2 f_{s,1}}{\partial z_2^2} = \frac{\partial^2 f_{s,1}}{\partial z_1 \partial z_2} = \frac{\partial^2 f_{s,1}}{\partial z_2 \partial z_3} = 0. \tag{C47}$$

Using eqs. (C40)-(C42) and (C46)-(C47), eq. (B24b) becomes,

$$\begin{aligned}
H_{{\rm w},s}(z) &= \frac{1}{2}\left[\left(\frac{z-\overline{q}_s}{\sigma_{\rm g}^2+\sigma_{\rm r,w}^2}\right)^2 \left(1-3\Phi_s(z)^2 r_{2,s}^{\rm R}(z)+2\Phi_s(z)^3 r_{3,s}^{\rm R}(z)\right) - \frac{1}{\sigma_{\rm g}^2+\sigma_{\rm r,w}^2}\left(1-\Phi_s(z)^2 r_{2,s}^{\rm R}(z)\right)\right] \\
H_{{\rm q},s}(z) &= \frac{1-d_s(z)}{d_s(z)}\left(\frac{z-\overline{q}_s}{\sigma_{\rm g}^2+\sigma_{\rm r,w}^2}\right)^2 \left(1-\Phi_s(z) r_{2,s}^{\rm R}(z)\right)\left(1-\Phi_s(z)^2 r_{2,s}^{\rm R}(z)\right) \\
H_{{\rm r},s}(z) &= -2(n-1) r_{2,s}(z)\frac{\gamma+(1-\gamma)(1-d_s(z))}{2\gamma+(1-\gamma)(1-d_s(z))}\Phi_s(z)^2\left(r_{2,s}^{\rm R}(z)-\Phi_s(z) r_{3,s}^{\rm R}(z)\right)\left(\frac{z-\overline{q}_s}{\sigma_{\rm g}^2+\sigma_{\rm r,w}^2}\right)^2.
\end{aligned} \tag{C48}$$

The coefficient of disruptive selection is then found by plugging eq. (C48) into eq. (B24) and using the different fecundity functions derived in section C.2.1 to compute the relevant quantities $r_{2,s}^{\rm R}(z)$ (eq. B6), $d_s(z)$ (eq. B7), $r_{2,s}(z)$ (eq. B8), $K_s(z)$ (eq. B16), $\Phi_s(z)$ (eq. B13), $r_{3,s}^{\rm R}(z)$ (eq. B21), $F_s(z)$ and $\overline{F}_s(z)$ (eq. C42).

The coefficient of disruptive selection is complicated but we can get further insights for the case where the population has converged to match the global average resource property, i.e. when $z^* = \overline{q}$ is convergence stable. In this case, relative fecundity reduces to

$$\frac{f_s(z)}{f(z)} = 1 \tag{C49}$$

for both states $s = 1, 2$ (using eqs. C35, C37 and C38). As a consequence, many relevant quantities that appear in eq. (C48) simplify. In particular, $d_s(z) = m$ and relatedness coefficients no longer depend on patch state, satisfying eqs. (C11) and (C22)-(C24). We thus obtain,

$$
\begin{aligned}
H_{\mathrm{w}}(z) &= \sum_{s\in\Omega}(1-\gamma)K_s(z)H_{\mathrm{w},s}(z)\pi_s \\
&= \frac{1}{2}\frac{1-\gamma}{\left(\sigma_{\mathrm{g}}^2+\sigma_{\mathrm{r,w}}^2\right)}\left[\frac{\sigma_{\mathrm{r,b}}^2}{\left(\sigma_{\mathrm{g}}^2+\sigma_{\mathrm{r,w}}^2\right)}\left(1-3(1-m)^2r_2^{\mathrm{R}}+2(1-m)^3r_3^{\mathrm{R}}\right)-\left(1-(1-m)^2r_2^{\mathrm{R}}\right)\right] \\
H_{\mathrm{q}}(z) &= \sum_{s\in\Omega}(1-\gamma)K_s(z)H_{\mathrm{q},s}(z)\pi_s \\
&= \frac{1-\gamma}{\left(\sigma_{\mathrm{g}}^2+\sigma_{\mathrm{r,w}}^2\right)}\frac{\sigma_{\mathrm{r,b}}^2}{\left(\sigma_{\mathrm{g}}^2+\sigma_{\mathrm{r,w}}^2\right)}\frac{1-m}{m}\left(1-(1-m)r_2^{\mathrm{R}}\right)\left(1-(1-m)^2r_2^{\mathrm{R}}\right) \\
H_{\mathrm{r}}(z) &= \sum_{s\in\Omega}(1-\gamma)K_s(z)H_{\mathrm{r},s}(z)\pi_s \\
&= -2\frac{1-\gamma}{\sigma_{\mathrm{g}}^2+\sigma_{\mathrm{r,w}}^2}(n-1)r_2\frac{\gamma+(1-\gamma)(1-m)}{2\gamma+(1-\gamma)(1-m)}(1-m)^2\left(r_2^{\mathrm{R}}-(1-m)r_3^{\mathrm{R}}\right)\frac{\sigma_{\mathrm{r,b}}^2}{\left(\sigma_{\mathrm{g}}^2+\sigma_{\mathrm{r,w}}^2\right)},
\end{aligned}
\tag{C50}
$$

for the three components of the coefficient of disruptive selection when $z = z^* = \overline{q}$. Summing these components yields,

$$
H(z) = H_{\mathrm{w}}(z)+H_{\mathrm{q}}(z)+H_{\mathrm{r}}(z) = \frac{1-\gamma}{\sigma_{\mathrm{g}}^2+\sigma_{\mathrm{r,w}}^2}\left(\chi_A\frac{\sigma_{\mathrm{r,b}}^2}{\sigma_{\mathrm{g}}^2+\sigma_{\mathrm{r,w}}^2}-\chi_C\right),
\tag{C51}
$$

for when $z = z^* = \overline{q}$ where

$$
\begin{aligned}
\chi_{\mathrm{A}} &= \left(\frac{1}{2}\left(1-3(1-m)^2r_2^{\mathrm{R}}+2(1-m)^3r_3^{\mathrm{R}}\right)+2(n-1)r_2\frac{\gamma+(1-\gamma)(1-m)}{2\gamma+(1-\gamma)(1-m)}(1-m)^2(r_2^{\mathrm{R}}-\left(1-m)r_3^{\mathrm{R}}\right)\right) \\
\chi_{\mathrm{C}} &= \frac{1}{2}\left(1-(1-m)^2r_2^{\mathrm{R}}\right).
\end{aligned}
\tag{C52}
$$

From eq. (C51), we readily get the branching condition (11) of the main text, i.e. we get that $H(z) > 0$ is equivalent to,

$$
\sigma_{\mathrm{r}}^2\left(\chi_{\mathrm{h}}E_{\mathrm{ST}}\ -\ (1-E_{\mathrm{ST}})\right)\ >\ \sigma_{\mathrm{g}}^2,
\tag{C53}
$$

where recall $\sigma_{\mathrm{r,b}}^2 = \sigma_{\mathrm{r}}^2E_{\mathrm{ST}}$, $\sigma_{\mathrm{r,w}}^2 = \sigma_{\mathrm{r}}^2(1-E_{\mathrm{ST}})$ and

$$
\chi_{\mathrm{h}} = \frac{\chi_{\mathrm{A}}}{\chi_{\mathrm{C}}}.
\tag{C54}
$$

###### C.2.4 Local competition

To investigate the impact of local competition when exploitation time is short, we assumed that density regulation additionally occurs before dispersal (between step 1) and 2) of the life cycle) so that the density of offspring within each patch is reduced to the same (large) offspring carrying capacity $C$ (where $C >> 1$). In this case, the philopatric and dispersal components of individual fitness (Appendix A.2) are now given by

$$\phi^{\mathrm{p}}_{s|s}(z_1, \boldsymbol{z}_{-1}, z) = (1-m)\frac{f_s(z_1,\boldsymbol{z}_{-1})}{\sum_{i=1}^{n} f_s(z_i,\boldsymbol{z}_{-i})}, \tag{C55}$$

$$\phi^{\mathrm{d}}_{s'|s}(z_1, \boldsymbol{z}_{-1}, z) = \pi_{s'} m\frac{f_s(z_1,\boldsymbol{z}_{-1})}{\sum_{i=1}^{n} f_s(z_i,\boldsymbol{z}_{-i})}, \tag{C56}$$

so that they reduce to

$$\begin{aligned} \phi^{\mathrm{d}}_{s'|s}(z) &= \pi_{s'}\frac{m}{n} \\ \phi^{\mathrm{p}}_{s|s}(z) &= \frac{1-m}{n} \end{aligned} \tag{C57}$$

in a monomorphic population.

We can then repeat our derivation of the selection gradient $S(z)$ (from eq. B3) and the disruptive selection coefficient $H(z)$ (from eq. B18) using the above. Note that in this case, we find that in terms of fecundity effects, the selection gradient and coefficient of disruptive selection can still be expressed respectively as eqs. (B10) and (B24) except that

$$\Phi_s(z) = 1. \tag{C58}$$

This is because the probability $\Phi_s(z)^2$ that two offspring born in the same patch of type $s$ compete with one another is equal to one with density regulation occurring before dispersal. In addition, since the same number of offspring is produced in each patch, we have

$$K_s(z) = 1 \tag{C59}$$

(as $f_s(z) = f(z)$ for all $s$).

**Selection gradient.** Using eq. (C58) and (C59), we obtain that the selection gradient is,

$$S(z) = \sum_{s\in\Omega} \pi_s(1-\gamma)\left(1-r_2^{\mathrm{R}}\right)\frac{\overline{q}_s - z}{\sigma^2_{\mathrm{g}}+\sigma^2_{\mathrm{r,w}}} = (1-\gamma)\left(1-r_2^{\mathrm{R}}\right)\frac{\overline{q}-z}{\sigma^2_{\mathrm{g}}+\sigma^2_{\mathrm{r,w}}}. \tag{C60}$$

Eq. (C60) shows that $z^* = \overline{q}$ is the only singular strategy, which is also convergence stable as,

$$\left.\frac{dS(z)}{dz}\right|_{z=z^*} = -(1-\gamma)\left(1-r_2^{\mathrm{R}}\right)\frac{1}{\sigma^2_{\mathrm{g}}+\sigma^2_{\mathrm{r,w}}} < 0. \tag{C61}$$

**Disruptive selection.** Similarly, we obtain that the three components of the coefficient of disruptive selection read as,

$$
\begin{aligned}
H_{\mathrm{w}}(z) &= \sum_{s\in\Omega}(1-\gamma)K_s(z)H_{\mathrm{w},s}(z)\pi_s \\
&= \frac{1}{2}\frac{1-\gamma}{\left(\sigma_{\mathrm{g}}^2+\sigma_{\mathrm{r,w}}^2\right)}\left[\frac{\sigma_{\mathrm{r,b}}^2}{\left(\sigma_{\mathrm{g}}^2+\sigma_{\mathrm{r,w}}^2\right)}\left(1-3r_2^{\mathrm{R}}+2r_3^{\mathrm{R}}\right)-\left(1-r_2^{\mathrm{R}}\right)\right] \\
H_{\mathrm{q}}(z) &= \sum_{s\in\Omega}(1-\gamma)K_s(z)H_{\mathrm{q},s}(z)\pi_s \\
&= \frac{1-\gamma}{\left(\sigma_{\mathrm{g}}^2+\sigma_{\mathrm{r,w}}^2\right)}\frac{\sigma_{\mathrm{r,b}}^2}{\left(\sigma_{\mathrm{g}}^2+\sigma_{\mathrm{r,w}}^2\right)}\frac{1-m}{m}\left(1-r_2^{\mathrm{R}}\right)^2 \\
H_{\mathrm{r}}(z) &= \sum_{s\in\Omega}(1-\gamma)K_s(z)H_{\mathrm{r},s}(z)\pi_s \\
&= -2\frac{1-\gamma}{\sigma_{\mathrm{g}}^2+\sigma_{\mathrm{r,w}}^2}(n-1)r_2\frac{\gamma+(1-\gamma)(1-m)}{2\gamma+(1-\gamma)(1-m)}\left(r_2^{\mathrm{R}}-r_3^{\mathrm{R}}\right)\frac{\sigma_{\mathrm{r,b}}^2}{\left(\sigma_{\mathrm{g}}^2+\sigma_{\mathrm{r,w}}^2\right)},
\end{aligned}
\tag{C62}
$$

where $z = z^* = \overline{q}$ so that

$$
H(z) = H_{\mathrm{w}}(z)+H_{\mathrm{q}}(z)+H_{\mathrm{r}}(z) = \frac{1-\gamma}{\sigma_{\mathrm{g}}^2+\sigma_{\mathrm{r,w}}^2}\left(\chi_A\frac{\sigma_{\mathrm{r,b}}^2}{\sigma_{\mathrm{g}}^2+\sigma_{\mathrm{r,w}}^2}-\chi_C\right), \tag{C63}
$$

where

$$
\begin{aligned}
\chi_A &= \left(\frac{1}{2}\left(1-3r_2^{\mathrm{R}}+2r_3^{\mathrm{R}}\right)+\frac{1-m}{m}\left(1-r_2^{\mathrm{R}}\right)^2-2(n-1)r_2\frac{\gamma+(1-\gamma)(1-m)}{2\gamma+(1-\gamma)(1-m)}(r_2^{\mathrm{R}}-r_3^{\mathrm{R}})\right) \\
\chi_C &= \frac{1}{2}\left(1-r_2^{\mathrm{R}}\right).
\end{aligned}
\tag{C64}
$$

From eq. (C63), it is straightforward to obtain that $H(z) > 0$ if and only if,

$$
\sigma_{\mathrm{r}}^2\left(\chi_1 E_{\mathrm{ST}}-(1-E_{\mathrm{ST}})\right) > \sigma_{\mathrm{g}}^2, \tag{C65}
$$

where

$$
\chi_1 = \frac{\chi_A}{\chi_C}. \tag{C66}
$$

is identical to $\chi_1$ where exploitation time is long (eq. C30). Hence, the condition for the emergence of polymorphism is identical to when exploitation time is long and resources vary only between patches (i.e. C65 is identical to C29 when $E_{\mathrm{ST}} = 1$).

### D Individual-based simulations

To accompany our mathematical analysis, we used *Nemo-Age* (Cotto et al., 2020) to simulate a diploid species of hermaphrodites subdivided among 2,000 patches of two types ($s = 1, 2$) with equal frequency ($\pi_1 = \pi_2 = 1/2$; see attached zip file for source code and makefile for compilation, to deposited in a repository upon acceptance). We fixed the global resource property to $\overline{q} = 50$. Then patches of type 1 and 2 were characterized by a normal distribution of resources with mean $\overline{q}_1 = \overline{q} + \sigma_{\mathrm{r,b}}$ and $\overline{q}_2 = \overline{q} - \sigma_{\mathrm{r,b}}$, respectively, and variance $\sigma^2_{\mathrm{r,w}}$ (the distribution was discretized into 51 equally sized bins to obtain $\pi_{j|s}$ for $j = 1,\ldots,51$ and $s = 1,2$). We explored various resource distribution by varying within- ($\sigma^2_{\mathrm{r,w}}$) and between-patch ($\sigma_{\mathrm{r,b}}$) variance to obtain five degrees of resource differentiation between patches ($E_{\mathrm{ST}}$=0, $E_{\mathrm{ST}}$=0.25, $E_{\mathrm{ST}}$=0.5, $E_{\mathrm{ST}}$=0.75 $E_{\mathrm{ST}}$=1) for two levels of the total resource variation ($\sigma^2_{\mathrm{r}}$=1 and $\sigma^2_{\mathrm{r}}$=2).

The life cycle in our simulations matched the one described in the main text (section 2.1) with the following events occurring each year: (1) Adults reproduced asexually making a number of offspring randomly drawn from a Poisson distribution with mean given by eq. (C1) (or eq. C34 depending on the scenario studied, and with each offspring a clonal copy of its hermaphroditic parent). (2) Offspring dispersed to a randomly chosen, non-natal patch with probability $m$ or remained philopatric with probability $1 - m$. (3) All the adults died (so we assumed $\gamma$=0 in all our simulations corresponding to a Wright-Fisher process). (4) $n = 10$ offspring were randomly sampled in each patch to become the adults of the next year.

To investigate the effects of sexual reproduction (and generate Figs. 6), we assumed that instead of step (1) above, hermaphrodite individuals mated randomly within patches. Specifically, we first picked the number of haploid eggs produced by each adult from a Poisson distribution with mean given by eq. (C1) (or eq. C34). Second, for each of these eggs, we picked an individual at random (with replacement) from the same patch to provide the fertilising haploid sperm (so that an individual self fertilises with probability $1/n$). The offspring individual resulted from the fusion of these two gametes.

Each individual $i$ expressed a consumer trait $z_i$ that controlled the feeding rate and determined individual fecundity $f_s(z_i, \boldsymbol{z}_{-i})$. The individual trait value was controlled by a single locus with additive allelic effects (so that an individual $i$ with alleles $a_{1,i}$ and $a_{2,i}$ expressed phenotype $z_i = a_{1,i} + a_{2,i}$; note that there is no environmental effect on phenotype in our simulations). Mutations occurred with probability $\mu = 0.00001$, with an effect whose size was picked from a normal distribution $\mathcal{N}(0, 0.05)$ (and added to the existing allelic effect following the continuum-of-allele model). In addition to the adaptive locus, each individual also carried an unlinked neutral locus that mutated according to the single-step mutational model (thought to capture microsatellite evolution): mutations occurred with probability $\mu = 0.00001$ and increased or decreased the allelic value by +1 or -1 (with reflective boundaries at 1 and 256).

We ran the simulations for 2,000 years, recording relevant summary statistics every 25 years, and storing the phenotypes of the entire population at year 2,000. Differentiation in allele frequencies at the neutral marker locus was computed with the *wc*-function of the *hierfstat* package following the Weir-Cockerham approach (Goudet, 2005). Differentiation in additive genetic effects of consumer traits ($Q_{\mathrm{ST}}$, which is identical to phenotypic differentiation $P_{\mathrm{ST}}$ in our simulations since we assumed no environmental effects) was calculated as follows. Phenotypic variance within and between patches was computed from an analysis of variance using the *aov*-function from the stats package in R (version 4.2.1, R Core Team, 2019). Additive genetic variance within populations was computed as $V_{\mathrm{G,w}} = MS_{\mathrm{within}}$ (the mean square within demes), the additive genetic variance between populations as $V_{\mathrm{G,b}} = (MS_{\mathrm{between}} - MS_{\mathrm{within}})/\eta_0$ with $\eta_0 = 10$ (where $MS_{\mathrm{between}}$ is the mean square between demes and $\eta_0$ is the average sample size per deme; e.g. see Storz et al., 2001; Martin et al., 2008). Then $Q_{\mathrm{ST}}$ was computed as $Q_{\mathrm{ST}} = V_{\mathrm{G,b}} / \left(V_{\mathrm{G,b}} + V_{\mathrm{G,w}}\right)$ under clonal reproduction, and $Q_{\mathrm{ST}} = V_{\mathrm{G,b}} / \left(V_{\mathrm{G,b}} + 2V_{\mathrm{G,w}}\right)$ under sexual reproduction. Note that in our model $V_{\mathrm{G,b}} = V_{\mathrm{P,b}}$ and $V_{\mathrm{G,w}} = V_{\mathrm{P,w}}$ as we do not have any environmental effects on phenotype expression.

### E Supplementary Figures

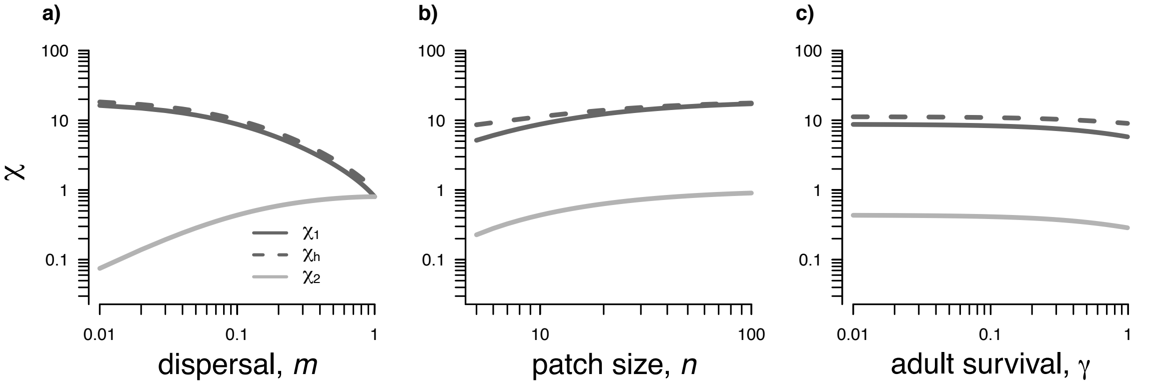


Figure S1: **The relative importance of local and spatial resource variation in promoting polymorphism.** The factors to spatial ($\chi_1$, $\chi_h$) and local ($\chi_2$) resource variation that appear in the branching conditions eq. (9) and (11) are plotted against dispersal $m$ (a), patch size $n$ (b), and adult survival $\gamma$ (c). If not specified otherwise, the dispersal probability was set to $m = 0.1$, the local patch size to $n = 10$, and adult survival to $\gamma = 0$. See eqs. (C28)-(C30) and eqs. (C54) in Appendix for details.

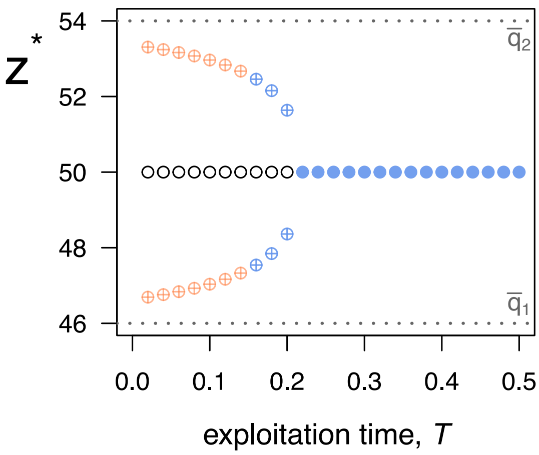


Figure S2: **Singular strategies and their stability under intermediate exploitation time $T$.** Bifurcation diagrams for the stability of singular strategies against exploitation time $T$ (with $\sigma_g^2 = 3$, $\gamma = 0$, $n = 10$, $m = 0.8$, $\sigma_{r,w}^2 = 1$, and $\sigma_{r,b}^2 = 16$). White circles indicate singular strategies that are evolutionary repellors; pink circles indicate singular strategies that are attractors and for which selection is stabilising (solid: $z^* = \overline{q}$; crossed: $z_1^* = \overline{q} - \theta$ or $z_2^* = \overline{q} + \theta$); blue circles indicate singular strategies that are attractors and for which selection is disruptive, i.e. evolutionary branching points (solid: $z^* = \overline{q}$; crossed: $z_1^* = \overline{q} - \theta$ or $z_2^* = \overline{q} + \theta$). Dotted lines indicate the average resource property in each habitat, $\overline{q}_1$ and $\overline{q}_2$. This shows that as exploitation time increases, polymorphism become more likely. This is because as exploitation time increases, interference competition also increases, favoring individuals to specialise on resources that are under less intense competition (eq. 6).

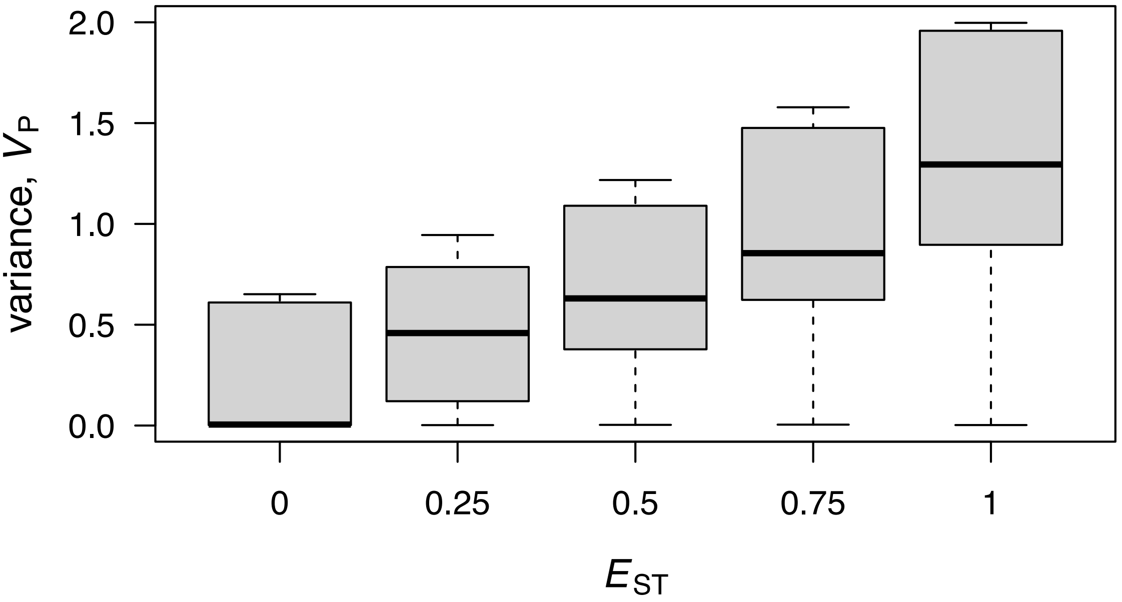


Figure S3: **Phenotypic variance according to resource differentiation among patches** $E_{ST}$. Box-plots for the distributions of phenotypic variance $V_P$ across multiple simulated populations after 2,000 years of evolution for different levels of $E_{ST}$. Parameters varied: $\sigma_r^2 = 1, 2$; $m = 0.05, 0.1, 0.2, \ldots, 1.0$; fixed parameters: $\sigma_g^2 = 1$, $n = 10$, $\gamma = 0$. These show that phenotypic variance increases with $E_{ST}$.
